# Inhibiting proBDNF to mature BDNF conversion leads to autism-like phenotypes *in vivo*

**DOI:** 10.1101/2020.06.12.149104

**Authors:** He You, Toshiyuki Mizui, Kazuyuki Kiyosue, Keizo Takao, Tsuyoshi Miyakawa, Koichi Kato, Mitsuru Otsuka, Ting Bai, Kun Xia, Bai Lu, Masami Kojima

**Author notes:** Correspondence should be addressed to M.K. and B.L.

## Abstract

Autism spectrum disorders (ASD) comprise a range of early age-onset neurodevelopment disorders with genetic heterogeneity. Most ASD related genes are involved in synaptic function, which is oppositely regulated by brain-derived neurotrophic factor (BDNF): the precursor proBDNF inhibits while mature BDNF (mBDNF) potentiates synapses. Here we generated a knock-in mouse line (BDNF^met/leu^) in which the conversion of proBDNF to mBDNF is inhibited. Biochemical experiments revealed residual mBDNF but excessive proBDNF in the brain. Similar to other ASD mouse models, the BDNF^met/leu^ mice showed decreased brain volumes, reduced dendritic arborization, altered spines, and impaired synaptic transmission and plasticity. They also exhibited ASD-like phenotypes, including stereotypical behaviors, deficits in social interaction, hyperactivity, and elevated stress response. Interestingly, the plasma level of proBDNF, but not mBDNF, was significantly elevated in ASD patients. These results suggest that proBDNF level, but not *Bdnf* gene, is associated with autism-spectrum behaviors, and identify a potential blood marker and therapeutic target for ASD.

## Introduction

Autism spectrum disorders (ASD) are early age-onset neurodevelopment disorders with genetic and clinical heterogeneity (Folstein & Rutter, 1977). There are three distinctive behavioral characteristics: social interaction deficits, communication disorders, and repetitive behaviors (De La Torre-Ubieta, Won, Stein, & Geschwind, 2016). A number of genes related to synaptic function have been genetically linked to ASD (De Rubeis et al., 2014), including *fmr1* in Fragile X syndrome (Bassell & Warren, 2008), *mecp2* in Rett syndrome (Amir et al., 1999), *scna1* in Dravet’s syndrome (Mahoney et al., 2009; Oakley, Kalume, & Catterall, 2011), and *cacna1* in Timothy syndrome (Splawski et al., 2004). Animal models with mutations of these genes display repetitive stereotypes, and deficits in social interaction similar to ASD patients (De La Torre-Ubieta et al., 2016). Interestingly, nearly all ASD animal models exhibit synaptic deficits, such as alterations in synaptic density, synaptic protein synthesis, and synaptic plasticity (De La Torre-Ubieta et al., 2016; Mullins, Fishell, & Tsien, 2016).

A key regulator for synaptic function is brain-derived neurotrophic factor (BDNF) (Barde, 1994; Ceni, Unsain, Zeinieh, & Barker, 2014; Greenberg, Xu, Lu, & Hempstead, 2009; Kaplan & Miller, 2007), a secretory protein best known for its roles in neuronal survival and differentiation during development, and synaptic function in the adult (Bibel & Barde, 2000; Park & Poo, 2013; Reichardt, 2006). Most of the ASD-related synaptic genes are up- or downstream of BDNF (Lu, Nagappan, Guan, Nathan, & Wren, 2013), although genetic studies have so far not provided a direct link between BDNF and ASD (De La Torre-Ubieta et al., 2016; De Rubeis et al., 2014; Miles, 2011). BDNF is initially synthesized as a precursor protein (proBDNF), which is proteolytic cleaved either intracellularly by furin and pro-protein convertases or extracellularly by plasmin and matrix metalloproteases (MMPs) to form mature BDNF (mBDNF) (Chao & Bothwell, 2002; Lee, Kermani, Teng, & Hempstead, 2001; Lu, Pang, & Woo, 2005b; Mowla et al., 2001; P. T. Pang et al., 2004; Woo et al., 2005).

A number of cell biological and physiological studies implicated that proBDNF is a secreted protein (Keifer, Sabirzhanov, Zheng, Li, & Clark, 2009; Nagappan et al., 2009; Woo et al., 2005; J. Yang et al., 2009). More importantly, proBDNF is not simply an inactive precursor, but an active protein that elicits biological functions through its own receptor (Lu et al., 2005b). There is good evidence that extracellular cleavage of proBDNF plays an important role in late phase long-term potentiation (LTP) and activity-dependent synaptic competition (Je et al., 2012; Je et al., 2013; Petti T. Pang, Nagappan, Guo, & Lu, 2016; P. T. Pang et al., 2004; F. Yang et al., 2009). Previous *in vitro* studies showed that proBDNF promotes the apoptosis of sympathetic neurons and cerebellar granule neurons, reduces the neurite outgrowth of basal forebrain cholinergic neurons (Teng et al., 2005; Volosin et al., 2006) and induces the retraction of dendritic spines of hippocampal neurons (H. Koshimizu et al., 2009). It has also been shown that proBDNF enhances long-term depression (LTD) in hippocampal slices (Liu et al., 2004; Massey et al., 2004; Rosch, Schweigreiter, Bonhoeffer, Barde, & Korte, 2005; Woo et al., 2005) These studies suggest that proBDNF, when secreted extracellularly, could elicit biological effects very different from and even opposite to mBDNF. Moreover, these apparently inhibitory effects are mediated by the pan-neurotrophin receptor p75^NTR^, but not by the high affinity BDNF receptor TrkB (H. Koshimizu et al., 2009; Teng et al., 2005; Volosin et al., 2006). Binding of proBDNF to the complex comprising p75^NTR^ and sortilin-related VPS10 domain containing receptor 2 (SorCS2) was shown to induce growth cone retraction by initiating the dissociation of the guanine nucleotide exchange factor Trio from the receptor complex (Deinhardt et al., 2011). Taken together, these *in vitro* studies have led to the “Yin-Yang” hypothesis that (i) BDNF and proBDNF elicit opposing biological functions and (ii) the conversion of proBDNF to mBDNF, or the ratio of proBDNF and mBDNF may determine the functional outcomes (Lu et al., 2005b).

A number of key questions remain: what is the physiological role of proBDNF? How important is the cleavage of proBDNF? A cleavage-resistant proBDNF knock-in mouse line was developed by mutating the two key arginine residues (Figure 1A, 127R and 128R) at the cleavage site of proBDNF (Yang et al., 2014). The homozygous mice were not viable, possibly due to a complete lack of mBDNF, in addition to the strong apoptotic effects of dramatically elevated levels of un-cleavable proBDNF in the brain. The heterozygous (*probdnf-HA/+*) mice, which have one copy of un-cleavable proBDNF, displayed a reduced dendritic complexity and impaired LTP similar to BDNF heterozygous mutant (BDNF+/-) (Yang et al., 2014). In addition, the *probdnf-HA/+* showed a reduced spine density, a decreased basal synaptic transmission, and an elevated LTD, which were not observed in BDNF+/-mice (Woo et al., 2005). Because the *probdnf-HA/+* mice expressed a high level of un-cleavable proBDNF but its mBDNF level was reduced by half, it is unclear whether the phenotypes observed were due to an increase in proBDNF or a decrease in mBDNF, or both. Moreover, the behaviors of the the *probdnf-HA/+* were not examined.

**Figure 1.**
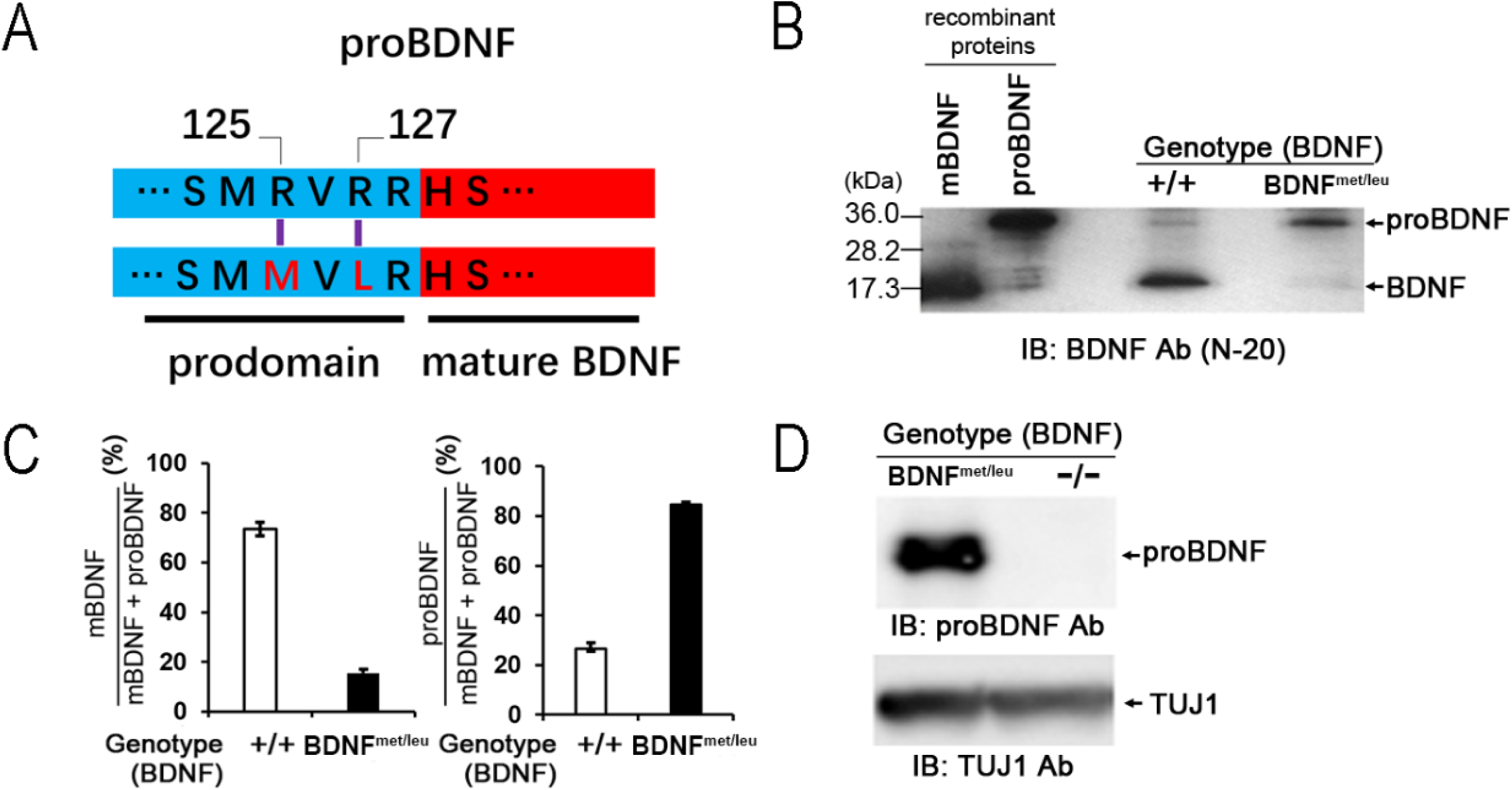
BDNFmet/leu knock-in mice expressing excessive proBDNF but residual mBDNF. **(A)** Schematic diagram showing the replacement of two arginine (R) with a methionine (M) and a leucine (L) in the conserved cleavage site of the *bdnf* gene to generate a cDNA encoding cleavage-resistant-type proBDNF. **(B)** Immunoblot analyses of BDNF protein expression in the hippocampus. Hippocampal lysates (30 μg of protein/lane) from 8-week-old littermates (wild type (+/+ or BDNF^+/+^) and BDNF^met/leu^ knock-in mice were processed for Western blot and probed with an antibody against the C-terminus of BDNF (N-20). Recombinant mature BDNF (mBDNF) and proBDNF proteins were used for comparison. **(C)** Quantification of the mBDNF or proBDNF bands on Western blot using Image J. N = 3 littermate mice per group. Note a dramatic increase in proBDNF and a dramatic decrease in mBDNF. In this and all other figures, bar graphs represented as the mean ± SEM. Student’s t-test was performed unless indicated otherwise. *: p<0.05; **: p<0.001. **(D)** Detection of proBDNF in BDNF^met/leu^ hippocampus, using a proBDNF-specific antibody. Hippocampal lysates from 10-day-old *Bdnf* homozygous knockout mice (-/-) were used as a negative control.

In the present study, we generated a proBDNF knock-in mouse line named BDNF^met/leu^, in which the arginine residues R125 and R127 at the cleavage site were converted to methionine (R125M) and leucine (R127L) (rs1048220 and rs1048221, respectively), based on previously reported human SNPs (H. Koshimizu et al., 2009). *In vitro* biochemical experiments demonstrated that these two mutations, either alone or together, markedly inhibit the cleavage of proBDNF (M. Koshimizu et al., 2015). The BDNF^met/leu^ mice expressed high levels of proBDNF but low levels of mBDNF, compared with their wild type (WT) littermates (H. Koshimizu et al., 2009). Unlike the homozygotes of proBDNF knock-in mice (*probdnf-HA/HA*) which died at birth, the homozygous BDNF^met/leu^ mice survived until adulthood (Kojima et al., 2020). This allowed us to perform detailed characterization of the adult BDNF^met/leu^ mice. Unexpectedly, this new line of mutant mice exhibited many morphological, physiological, and behavioral phenotypes often seen in ASD patients. Further, preliminary characterization of ASD patients revealed a significant increase in the plasma level of proBDNF. Our results point to a possible non-genetic mechanism of ASD and suggest a blood marker potentially useful in the clinic.

## Results

### Generation of BDNF^met/leu^ mice predominantly expressing proBDNF

We generated a knock-in mouse line in which the endogenous *Bdnf* allele was replaced with proBDNF containing human SNPs that changed two arginines proximal to the cleavage site to methionine (R125M) and leucine (R127L), respectively (Figure 1A). We previously reported that these amino acid changes led to inefficient conversion of proBDNF to mature BDNF in cultured neurons (H. Koshimizu et al., 2009). Immunoblot analyses using a polyclonal BDNF antibody (N-20, Santa Cruz) demonstrated that hippocampal lysates from 8-week-old homozygous mutant (BDNF^met/leu^) mice contained an excess amount of proBDNF and only a residual amount of mBDNF, whereas those from wild-type (WT) littermates (or BDNF^+/+^) contained more mBDNF than proBDNF (Figure 1B). This opposite ratio of mBDNF and proBDNF was quantitatively confirmed by Western blot analysis using a monoclonal anti-pan-BDNF antibody (clone; 3C11, Icosagen) (Kojima et al., 2020) (Figure 1C). A band with the same size as proBDNF was also observed via immunoblotting using another previously-generated proBDNF-specific antibody (Nagappan et al., 2009), further validating that this was indeed proBDNF (Figure 1D). Taken together, we generated a proBDNF knock-in mouse line that expresses a high level of proBDNF with a residual amount of mBDNF.

Despite insufficiency in the proBDNF cleavage, the BDNF^met/leu^ homozygous mice were born in a Mendelian manner, and approximately 95% of the homozygous mutant mice survived to adulthood (Kojima et al., 2020). We therefore performed all subsequent experiments using BDNF^met/leu^ homozygous, but not heterozygous, mice. The survival of the BDNF^met/leu^ mice to adulthood was unexpected because the homozygous proBDNF knock-in mice with different mutations at the proBDNF cleavage site (RVRR→RVAA) died shortly after birth, as reported previously (Yang et al., 2014). Unlike the proBDNF knock-in mice, the morphology and TrkB activation of the hearts of BDNF^met/leu^ mice were normal (Kojima et al., 2020). These morphological and immunohistochemical results suggest that the inefficient conversion of proBDNF into mBDNF due to R125M/ R127L mutation may not influence the survival of animals and/or development of the heart.

### Brain volumes and dendritic complexity in BDNF^met/leu^ mice

The brains of the BDNF^met/leu^ mice were smaller than their WT counterparts (Figure 2A). The mean wet weight of the BDNF^met/leu^ brains was 15.1 ± 1.3% lower than that of the BDNF^+/+^ brains (Figure 2B). Cavalieri analysis of Nissl-stained sections revealed that the whole brain volume of the BDNF^met/leu^ group was 17.7 ± 1.56% lower than that of the BDNF^+/+^ group (Figure 2C). Similarly, the volume of the BDNF^met/leu^ hippocampus and cortex was reduced by 14.0 ± 2.3% and 15.8 ± 1.5%, whereas the reduction in olfactory bulb was not significant, compared with the BDNF^+/+^group (Figure 2C). Thus, inefficient cleavage of proBDNF affects the volume of the adult brain, particularly in the hippocampus and cerebral cortex, two areas known to express high levels of BDNF mRNA (Timmusk et al., 1993).

**Figure 2.**
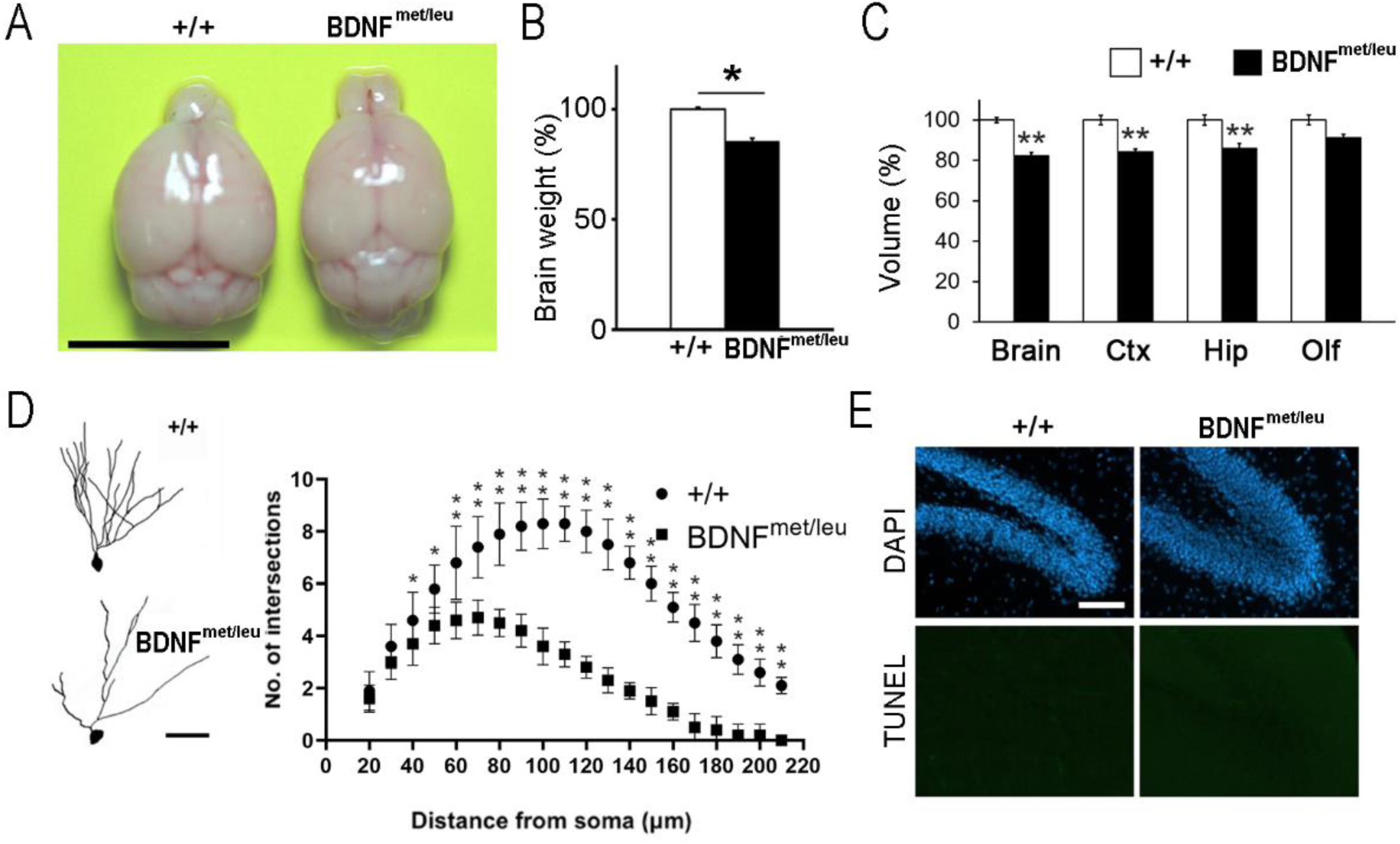
Decrease in brain volume and dendritic arborization in BDNFmet/leu mutants. **(A)** Gross view of representative brains of +/+ and BDNF^met/leu^ littermates. Scale bars: 10 mm. **(B)** Quantification of brain weights of 2-month-old mice with the indicated genotype. N = 6 animals per group. **(C)** Cavalieri analyses of Nissl-stained brain sections of +/+ and BDNF^met/leu^ littermates showing volumes of whole brain and its sub-regions. N = 6 animals per genotype. Ctx: cortex; Hip: hippocampus; Olf: olfactory bulb. **(D)** Sholl analyses of dentate gyrus (DG) neurons from 8-week-old mice N = 10 neurons per group. Left: an example of the Golgi-stained neurons. Scale bars: 50 μm. **(E)** Detection of apoptotic cells in DG granule cell layers. Coronal sections of hippocampi from 8-week-old mice were stained using a TUNEL assay. Scale bars: 100 μm.

Since proBDNF is highly expressed in dentate gyrus (DG) (Zhou et al., 2004) the dendritic complexity of the DG neurons was analyzed using Golgi staining. The BDNF^met/leu^ homozygous mice had a marked decrease in dendritic arbor complexity compared with their BDNF^+/+^ littermates (Figure 2D, left panel). The difference became evident at 70 μm from cell body (Figure 2D, right panel). Nevertheless, there was no significant difference between the diameters of the cell bodies in the BDNF^met/leu^ and WT animals (data not shown). No apparent apoptotic cells were detected in the DG regions of brains from each group (Figure 2E). Thus, inhibition of proBDNF cleavage may lead to impairment of dendritic complexity but not neuronal survival at least in hippocampal DG region in animals.

The reduction in brain volume was also reported in mouse models with autism-like phenotypes (De La Torre-Ubieta et al., 2016), such as Nlgn3 KO (Baudouin et al., 2012), Nlgn4 KO (Jamain et al., 2008) and ARX cKO mice (Fulp et al., 2008). Furthermore, reduced dendritic complexity was observed in models with changes in the expression of ASD related proteins, such as decreased TAO2 (de Anda et al., 2012) or elevated UBE3A (Khatri et al., 2018). Thus, the BDNF^met/leu^ mice manifest morphological changes similar to other ASD models.

### Dendritic spine changes in BDNF^met/leu^ mice

Next, we performed a detailed analysis of the dendritic spine morphologies and density in BDNF^met/leu^ hippocampus. The maximal length, maximum width, and density of spine protrusions in the secondary dendritic segments of hippocampal pyramidal neurons in the stratum radiatum of the CA1 area were determined (Figure 3A). We found no significant difference in the mean protrusion lengths between the BDNF^met/leu^ and WT groups (*P* = 0.25) (Figure 3B). However, the mean protrusion width of the BDNF^met/leu^ mice was significantly smaller than that of the WT mice (Figure 3C).

**Figure 3.**
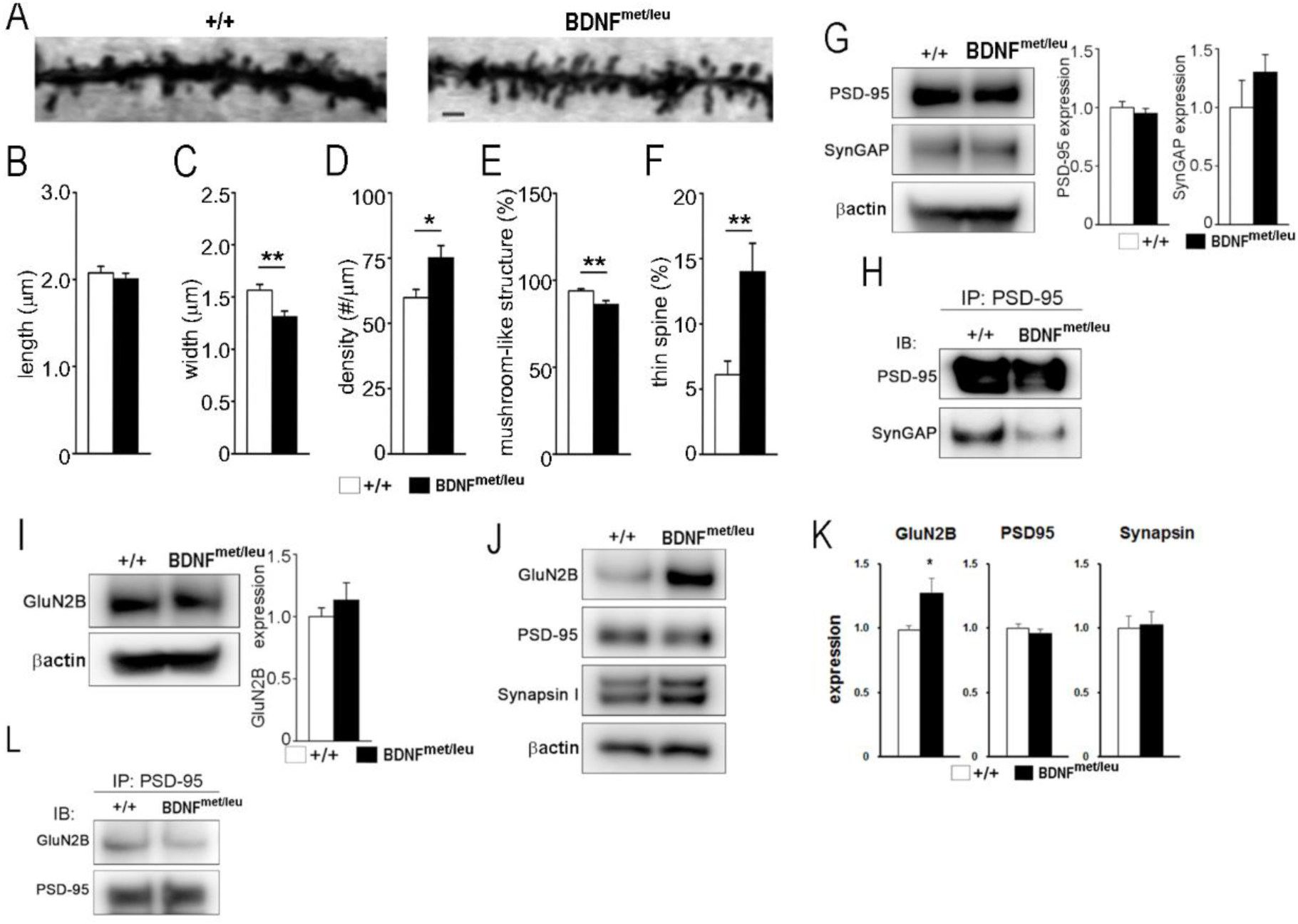
Alterations in dendritic spines and interactions of postsynaptic proteins. **(A)** Representative high-magnification images of Golgi-stained CA1 pyramidal neurons isolated from 9-week-old +/+ and BDNF^met/leu^ littermates. Scale bar, 2 μm. **(B–F)** Quantitative analyses of the morphologies of secondary dendritic segments of hippocampal pyramidal neurons in the stratum radiatum of the CA1 area. The length (B), width (C), and density (D) of the spine protrusions, as well as the percentages of mushroom-like (E) and thin (F) spines, were quantified. Data are represented as the mean ± SEM of N = 3 mice per genotype and n = 33 (+/+) or n = 36 (BDNF^met/leu^) neurons per mouse. **(G)** Immunoblot analyses of PSD-95 and SynGAP expression levels in hippocampal lysates (25 μg protein) from +/+ and BDNF^met/leu^ mice. The expression levels of PSD-95 and SynGAP were normalized to that of β-actin. The left panel shows a representative blot and the right panel shows quantification of the data (PSD-95, p = 0.19; and SynGAP, p = 0.73, by Student’s t-tests). N = 3 mice per group. **(H)** The interaction of PSD-95 with SynGAP in +/+ and BDNF^met/leu^ hippocampus. Hippocampal lysates were immunoprecipitated with an anti-PSD-95 antibody, followed by immunoblotting using anti-PSD-95 and anti-SynGap antibodies. Note a selective decrease in SynGap in BDNF^met/leu^ hippocampus. **(I)** Immunoblot analyses of GluN2B expression in hippocampal lysates from +/+ and BDNF^met/leu^ mice. The expression levels of GluN2B were normalized to that of β-actin. The left panel shows a representative blot and the right panel shows quantification of the data (N = 3 mice per group. p = 0.18 by Student’s t-test). **(J-K)** The level of synaptic proteins in hippocampal synaptosomal preparations from +/+ and BDNF^met/leu^ mice. Note a selective increase in GluN2B in BDNF^met/leu^ hippocampus. **(L)** The interaction of GluN2B with PSD-95 in +/+ and BDNF^met/leu^ hippocampi. Hippocampal lysates (2 mg protein) were immunoprecipitated (IP) with an anti-PSD-95 antibody, and equal aliquots of the immunoprecipitants were subjected to immunoblotting (IB) with the indicated antibodies. Note that the interaction between PSD95 and GluN2B was decreased in BDNF^met/leu^ hippocampus, compared with that in the +/+ hippocampus.

A previous report using heterozygous *probdnf-HA/+* mice showed a marked decrease in total spine density, without detailed classification of different types of spines (Yang et al., 2014). It was also unclear whether the decrease in spine density was due to an increase in proBDNF or a decrease in mBDNF. In contrast, we observed a significant increase in total spine density (Figure 3D). According to previous studies, the spine protrusions could be classified as thin (a protrusion with length to width ratio >2) or mushroom-like ones (Bourne & Harris, 2008; Harris, Jensen, & Tsao, 1992; Matsuzaki et al., 2001). Detailed quantitative analyses of CA1 neuron dendrites revealed a small but significant decrease in mushroom-like spines (Figure 3E), but a more than 2-fold increase in thin spines (Figure 3F), in the BDNF^met/leu^ mice. These results reveal that our BDNF^met/leu^ mice are distinct from the previously reported *probdnf-HA/+* mice and suggest that the prevention of proBDNF cleavage results in instability of dendritic spines in hippocampus. Selective increase in the density of thin spines (or filopodia) has been observed in Fmr1 KO mice (de Vrij et al., 2008), a mouse model for Fragile X Syndrome (Bernardet & Crusio, 2006; Gross et al., 2015; Huber, Gallagher, Warren, & Bear, 2002; Lim et al., 2014). Selective decrease in the density of mushroom-like dendritic spines has been reported in Tg1 mice that over-express human MeCP2, a mouse model for Rett syndrome.

### Altered postsynaptic scaffold

Recent studies suggest that some of the biological effects of proBDNF require its interaction with p75^NTR^ and the sortilin family proteins such as SorCS2 and SorCS3 (Glerup et al., 2016). SorCS3 is expressed at a high level in the hippocampal CA1 region and is localized to the postsynaptic density (PSD), and the loss of SorCS3 in mice leads to the impairment of hippocampal LTD (Breiderhoff et al., 2013). We therefore investigated whether the proBDNF signaling machinery is intact in the BDNF^met/leu^ hippocampus. We found that the expression levels of p75^NTR^, SorCS2, and SorCS3 in the hippocampus were not affected by the BDNF^met/leu^ mutation (Figure 3-supplement 1). Thus, proBDNF signaling through the p75^NTR^-SorCS2 and/or SorCS3 receptor complex occurs normally in the BDNF^met/leu^ mice.

The deficits in postsynaptic scaffold proteins have been reported in several ASD mouse models (Mullins et al., 2016; Peça et al., 2011). Two proteins have been implicated in spine structure/function: PSD-95, a postsynaptic scaffold protein abundant in spines, and SynGAP, a Ras GTPase-activating protein known to regulate spine morphology (Kim, Liao, Lau, & Huganir, 1998; Vazquez, Chen, Sokolova, Knuesel, & Kennedy, 2004). The expression levels of PSD-95 and SynGAP were not affected by the BDNF^met/leu^ mutation (Figure 3G). However, immunoprecipitation studies revealed that the interaction of PSD-95 and SynGAP was attenuated markedly in the lysates prepared from 9-week-old BDNF^met/leu^ mice, compared with the WT control (Figure 3H).

proBDNF has been shown to promote hippocampal LTD by regulating the expression of the GluN2B subunit of the N-methyl-D-aspartate receptor (NMDAR)(Woo et al., 2005). The expression of hippocampal GluN2B were comparable between WT and BDNF^met/leu^ animals (Figure 3I). Interestingly, the level of GluN2B was significantly higher in the crude synaptosomal fraction (P2) prepared from BDNF^met/leu^ hippocampi (Figure 3J and K). In contrast, its interaction with PSD-95 was slightly decreased (Figure 3L). However, the levels of PSD-95, synapsin I, or beta-actin in the same P2 fraction were not altered (Figure 3J and K). Overall, these data suggest that proBDNF signaling *in vivo* leads to a down-regulation of the molecular interaction of scaffolding proteins in dendritic spines but an up-regulation of GluN2B expression at extrasynaptic sites in the hippocampal neurons. These findings are reminiscent to the impairments in NMDA receptor function observed in the Shank3 mutant (Duffney et al., 2015) and Shank2 exons 6-7 KO mice (Won et al., 2012), which are also used as ASD models..

### Impaired synaptic transmission and plasticity

We further examined whether deficits in dendritic spines would alter synaptic function in Schaffer collateral-CA1 pyramidal neurons (Mizui et al., 2015). The slope of the input/output (I/O) curve was significantly reduced, suggesting an impairment in basal synaptic function in BDNF^met/leu^ synapses (Figure 4A). However, paired-pulse facilitation (PPF) was not altered in CA1 synapses of the BDNF^met/leu^ mice, indicating normal presynaptic function (Figure 4B). The decrease in mushroom spines but normal PPF suggest that the reduction in basal synaptic transmission could be mediated by post-synaptic rather than pre-synaptic mechanisms.

**Figure 4.**
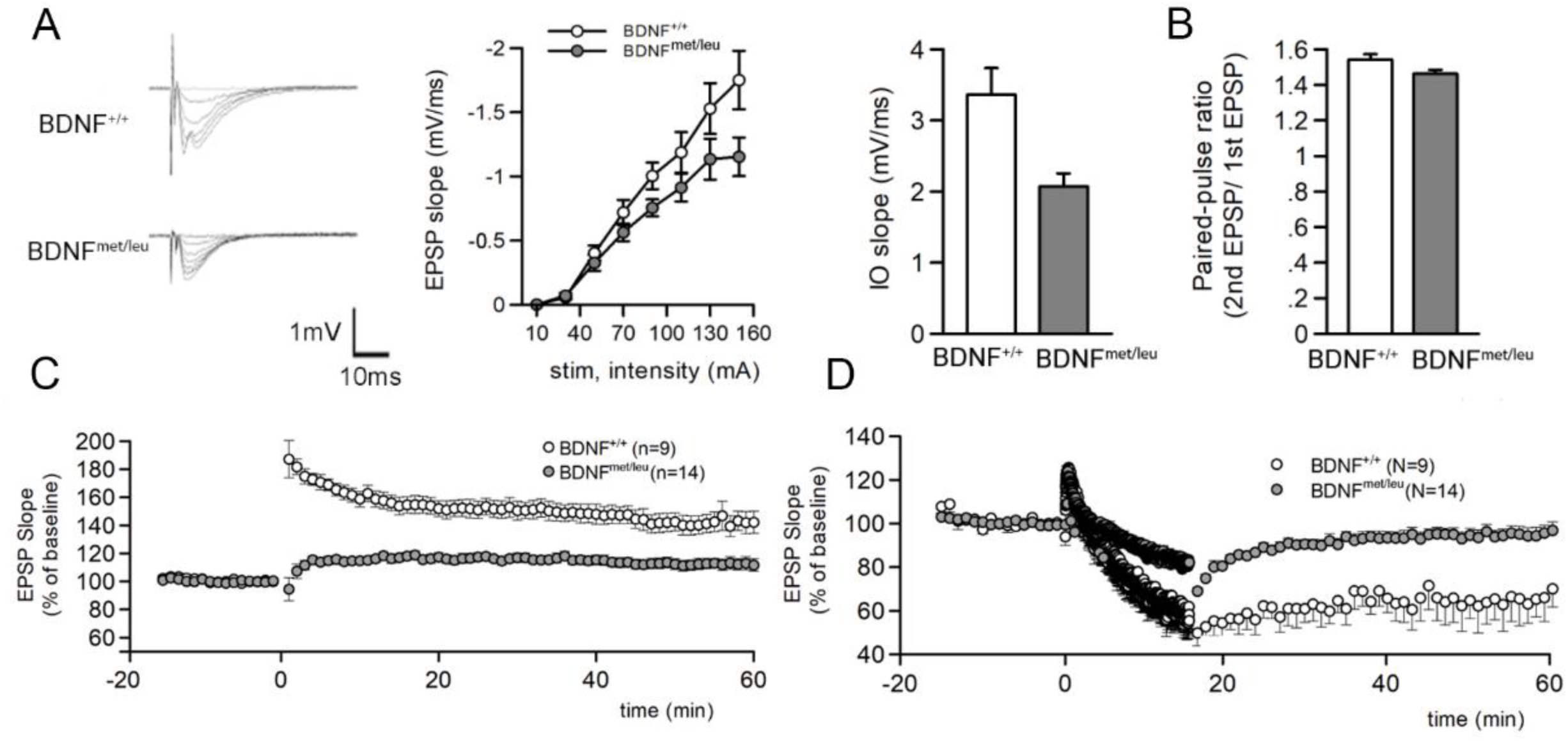
Impairments in synaptic transmission and plasticity. (**A**) A decrease in basal synaptic transmission in BDNF^met/leu^ mice. Representative traces (Left), quantitative analysis (Middle), and slope (right) of the input-output (I/O) curve of field excitatory postsynaptic potentials (fEPSPs) recorded in Schaffer collateral-CA1 pyramidal neurons in the hippocampus of +/+ and BDNF^met/leu^ mice. The slopes of fEPSP were plotted against the fiber volley amplitude. +/+: N= 4 animals, n=9 slices; BDNF^met/leu^: N= 4 animals, n=9 slices. (**B**) Normal paired-pulse facilitation (PPF) in BDNF^met/leu^ mice. Two stimuli were applied at various inter-pulse intervals (IPI, 10 to 100 msec). The ratios of the second and first fEPSP slopes were calculated, and mean values were plotted against different inter-pulse intervals (IPI, 10 to 100 msec). +/+: N= 4 animals, n=9 slices; BDNF^met/leu^: N= 4 animals, n=9 slices. (**C**) Impairments of long-term potentiation (LTP) in BNDF^met/leu^ mice. The fEPSP slopes were plotted against time (in minutes) before and after theta-burst stimulation (TBS, 5 pulses at 100Hz per burst, 10 bursts with an interval of 200 msec). n in this and (D) indicates number of slices used. (**D**) Impairments of long-term depression (LTD) in BNDF^met/leu^ mice. The experiments were carried out in the same way as above except low-frequency stimulation (1Hz 900 pulse) were applied to hippocampal slices to induce LTD.

We next examined long-term synaptic plasticity in BDNF^met/leu^ mice. Previous studies have shown that the application of exogenous proBDNF enhanced LTD in juvenile (5-week-old) hippocampal slices (Woo et al., 2005). LTD was elevated in 3-week-old hippocampal slices from the previously reported in the heterozygous *probdnf-HA/+* mice, which have half of its mBDNF but elevated proBDNF (Yang et al., 2014). We tested LTD in hippocampal slices from 3-week-old BDNF^met/leu^ mice which predominantly express proBDNF with residual mBDNF (Figure 1C). The Low-frequency stimulation (LFS; 1 Hz, 900 pulses, 15 min) was applied to Schaffer collaterals of hippocampal slices. In marked contrast to what was observed in *probdnf-HA/+* mice, both induction and expression of LTD was severely impaired in BDNF^met/leu^ mice (Figure 4D). We further examined whether hippocampal LTP was altered in BDNF^met/leu^ mice. Two-month-old hippocampal slices were used, field EPSP slopes at CA1 synapses were recorded, and high-frequency stimulation (HFS, 100Hz, 1 second) was applied. Although control mice expressed normal LTP (Figure 4C), the BDNF^met/leu^ mice failed to express LTP (Figure 4C). Taken together, it appears that BDNF^met/leu^ mice, similar to most ASD models (De La Torre-Ubieta et al., 2016), exhibit deficits in all aspects of synaptic functions, including synaptic transmission and plasticity. For example, the Shank2 exons 6–7 KO mice showed impairment in both LTD and LTP (Won et al., 2012).

### Stereotypes and deficits in social interactions

Reduced brain volume, decreased dendritic complexity, increased thin spines, as well as deficits in postsynaptic scaffold proteins and synaptic functions are all characteristics of mouse models of ASD. In humans, ASD is a group of developmental brain disorders with several distinctive features, including reduced social interactions, language problems, and repetitive and restrictive behaviors (De La Torre-Ubieta et al., 2016; Mullins et al., 2016). While it is difficult to investigate language problems in animals, several mouse ASD models display common autistic behaviors such as hyperactivity, repetitive and stereotyped behaviors, and anxiety (De La Torre-Ubieta et al., 2016). We therefore investigate whether BDNF^met/leu^ mice have some of the ASD-related behavioral phenotypes.

An obvious repetitive and stereotypical behavior seen in the BDNF^met/leu^ mice was “stargazing.” In home cages, BDNF^met/leu^ mice displayed repeated head-tossing very similar to that seen in the classic “stargazer” mouse (Figure 5A, Movie#1) (L. Chen et al., 2000). Quantitative analyses showed that the mutant mice underwent stargazing over 50 times per min (Figure 5B). Such behavior was hardly seen in WT mice.

**Figure 5.**
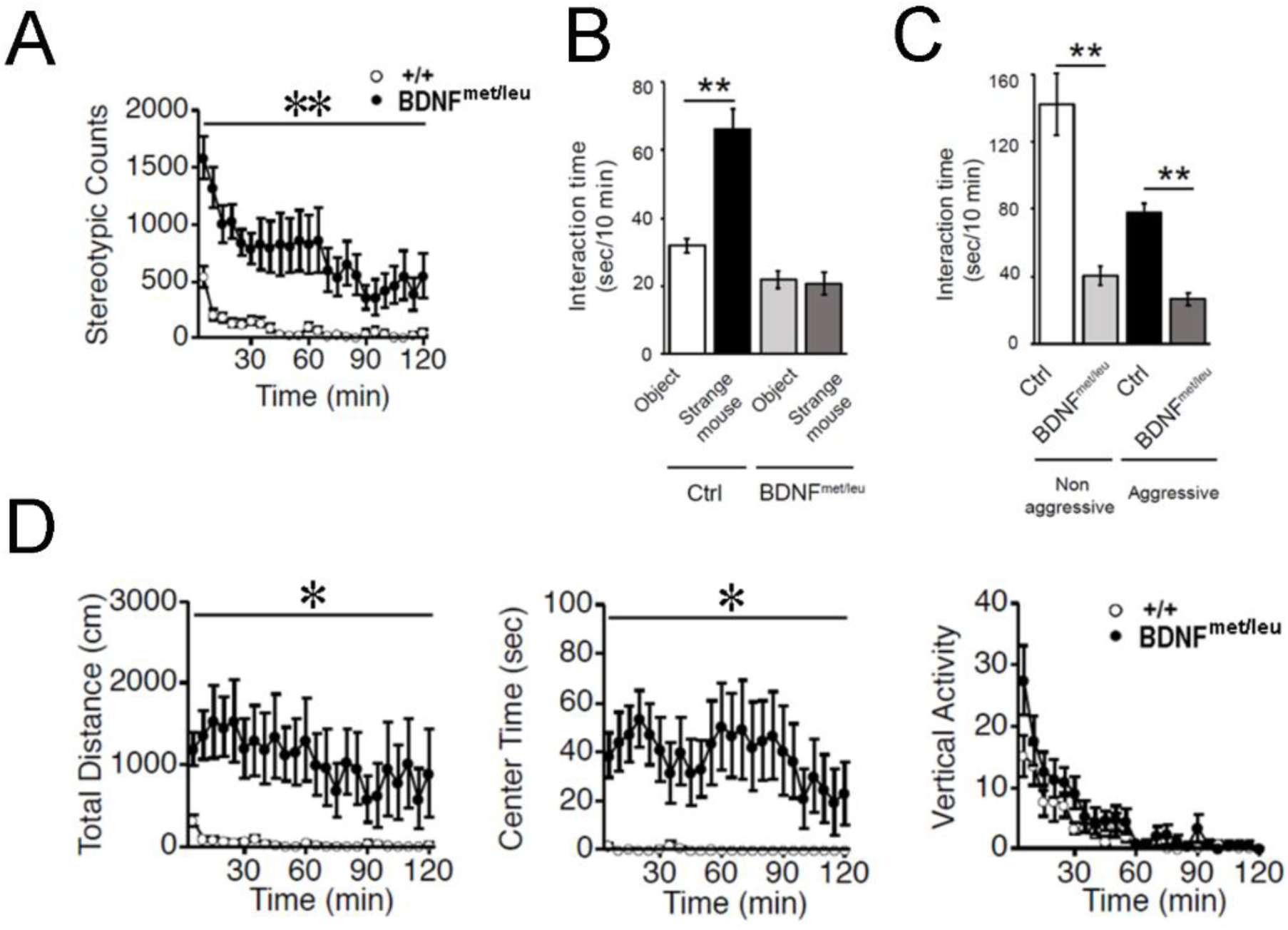
Autism spectrum disorders (ASD)-related behavioral phenotypes. (**A**) Stereotype behavior. In the same open field, the number of “star-gazing” like behaviors were counted over a 120-min period. (**B**) Social-interaction test in the open field. Interaction time was measured as the time mice spent in the quadrant of target object or stranger mouse within in a 10-min period. When presented both an object and a stranger mouse simultaneously, +/+ mice spent more time in the quadrant to interact with the caged stranger mouse. However, BDNF^met/leu^ mice had no preference for the stranger mouse. **(C)** Social interactions of freely moving BDNF^met/leu^ and +/+ littermates with aggressive or non-aggressive test mice. A mouse generally interacts less with an aggressive littermate than with a non-aggressive one. In contrast, BDNF^met/leu^ mice displayed a decreased time in both non-aggressive and aggressive interactions. N = 7 mice per genotype. **P < 0.01. (**D**) BDNF^met/leu^ mice exhibit severe hyperactivity. Mice were placed in a standard open field, and total distance traveled (left), the amount of time spent in the center of the open field (center), and vertical activity (right) were counted in a 120-min period.

Next, we performed two tests to assess social interaction by BDNF^met/leu^ mice. First, a three-chamber test was performed to examine whether a BDNF^met/leu^ mouse placed in the middle chamber prefers a stranger mouse (in one side chamber) to a novel object (in the opposite side chamber). The WT mice spent twice more time with a mouse than with an object (Figure 5B). In contrast, the BDNF^met/leu^ mice showed no preference to social partner, spending equal amount of time interacting with the mouse partner and the object (Figure 5B). Second, we measured the interaction of a BDNF^met/leu^ mouse with their littermates in the home cage. A mouse generally interacts less with an aggressive littermate than with a non-aggressive one. We therefore measured the interactions with the two types of littermates separately. BDNF^met/leu^ mice displayed dramatically reduced interaction time with both aggressive and non-aggressive littermates, compared with the WT mice (Figure 5C). These results indicated that social interaction was impaired in BDNF^met/leu^ mice.

### Hyperactivity and elevated stress response

As most ASD mice models are reportedly hyperactive (De La Torre-Ubieta et al., 2016), we tested locomotion and motor ability of BDNF^met/leu^ mice. In an open field test, the BDNF^met/leu^ mice traveled much longer distances, approximately 1000cm/min, than their WT littermates (Figure 5D, left). They also traveled mostly in the center of the field, spending approximately 40sec/min there (Figure 5D, center). There was no difference in their vertical activities between the BDNF^met/leu^ mice and their WT littermates (Figure 5D, right). Interestingly, the hyperactivity phenotype appeared to be more apparent during the nighttime when the mice are mostly awake (Figure 5-supplement 1).

While hyperactive, ASD patients often become restrictive and immobile when challenged or stressed (De La Torre-Ubieta et al., 2016). Tail-suspension test (TST) is often used to test stress-induced immobile behavior in mice. The BDNF^met/leu^ mice (10-12-week old) showed significantly longer immobility time in TST than their WT littermates (Figure 6A). However, unlike in WTs, the immobility time was not shortened by fluoxetine (20 mg/kg, 30 min, i.p.), a selective serotonin reuptake inhibitor commonly used as an antidepressant (Crowley, Blendy, & Lucki, 2005)(Figure 6A). To determine whether prolonged immobility in TST is associated with stress, we determined blood concentrations of corticosterone, an adrenal steroid known to increase in response to stress. Blood samples were collected from the tail vein of BDNF^met/leu^ and WT littermates reared in the same cage for 2 weeks. As expected, the blood corticosterone concentration in resting time in WT mice was comparable to that reported previously (Dugovic, Maccari, Weibel, Turek, & Van Reeth, 1999). In contrast, the BDNF^met/leu^ mice demonstrated a 3-fold higher blood corticosterone concentration (Figure 6B, Before stress), suggesting that BDNF^met/leu^ animals are severely stressed compared with WT littermates reared in the same cage. However, 60 min after the exposure to immobilization stress, both BDNF^met/leu^ and WT mice exhibited similar, elevated levels of corticosterone (Dugovic et al., 1999), (Figure 6B, right, After stress). Consistent with these results, in an unstressed home cage environment, the net weight of the adrenal glands, which secrete corticosterone (Jankord & Herman, 2008), was significantly higher in BDNF^met/leu^ than that in WT mice (Figure 6C, Adrenal gland/Body). As a negative control, the weight of the kidney did not differ between the BDNF^met/leu^ and WT animals ((Figure 6C, Kidney/Body). These results suggest that mice deficient in proBDNF-processing are sensitive to stress and exhibit activation of the HPA axis under normal conditions.

**Figure 6.**
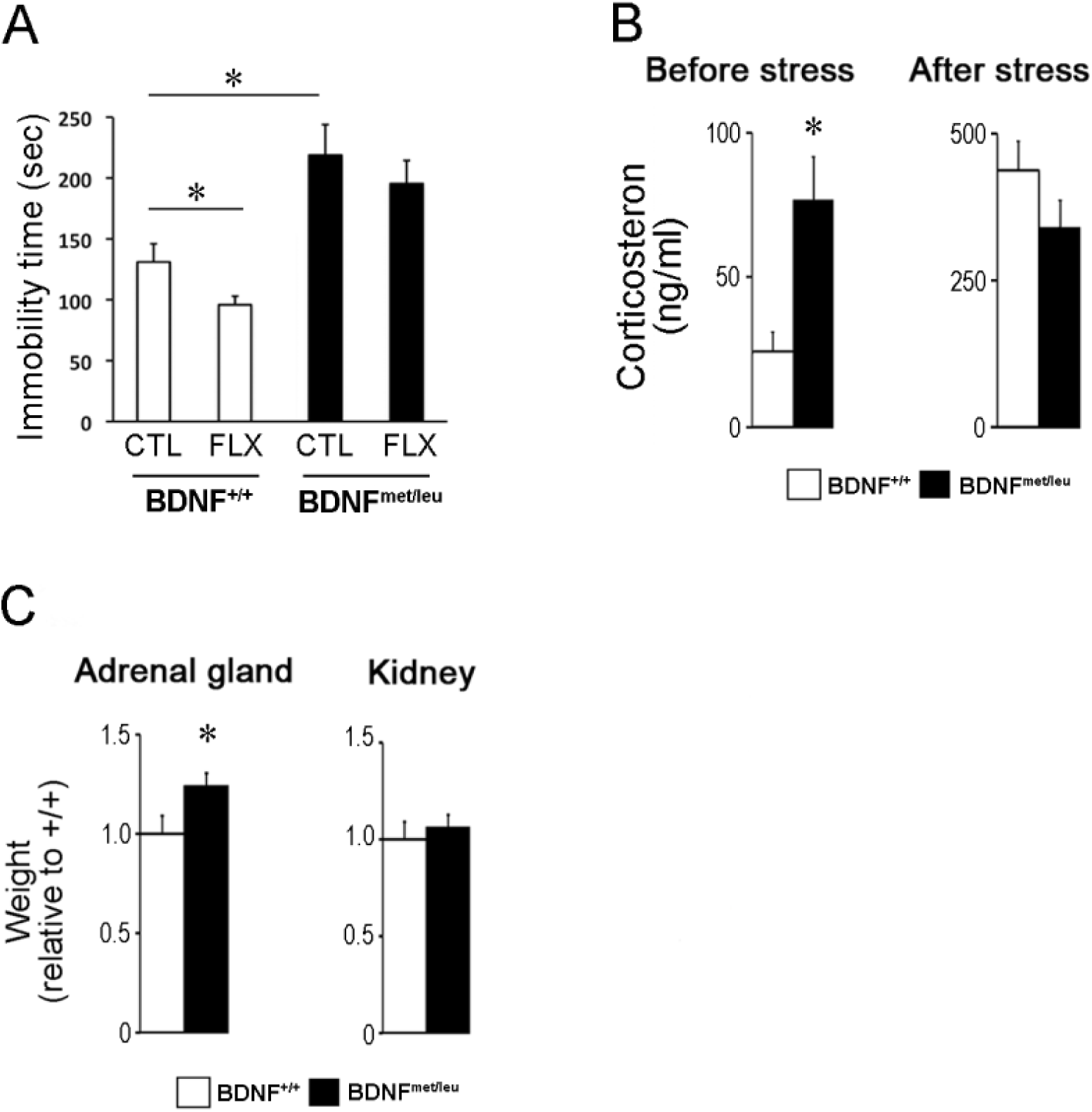
Elevated stress response in BDNFmet/leu mice. **(A)** Tail suspension test of BDNF^+/+^ and BDNF^met/leu^ mice at the age of 10∼12 weeks. Animals were administered saline or fluoxetine 30 min before the tail suspension test (10 mg/kg i.p.). *, p < 0.05; ANOVA with post-hoc test. N=7∼8 littermate animals. **(B)** Corticosterone levels in the serum before and after acute stress exposure. The tail blood samples were collected in group-housed BDNF^+/+^ and BDNF^met/leu^ mice (10-week-old, N = 8 per genotype) (Left panel, before stress). After an additional 2 weeks of group-housing, the animals were subjected to immobilization stress (60 min) and the tail blood samples were collected immediately (Right panel, after stress). The serum corticosterone concentration was determined immediately after sampling. **(C)** The relative weights of the adrenal gland and kidney in 8-week-old littermate animals (N = 6 per genotype) with the indicated genotypes. The organ weights were normalized to the body weight of each animal.

### Plasma proBDNF and BDNF levels in ASD patients and controls

A number of previous studies have suggested that blood (serum) level of BDNF is proportional to its level in the brain (Klein et al., 2011). Given that the BDNF ^met/leu^ mice expressed excessive proBDNF in the brain, we determined whether blood proBDNF concentration was also elevated. However, blood platelets contain a large quantity of BDNF (Fujimura et al., 2002; Yamamoto & Gurney, 1990), which could interfere the accurate measurement of BDNF in the blood. Furthermore, the majority of ELISAs used in previous literature could not distinguish between proBDNF and mBDNF. We therefore measured proBDNF specifically, using plasma, which do not contain platelets or its content instead of serum. We used a sandwich ELISA, which included a capture antibody against the C-terminal BDNF, a detection antibody against the pro-domain of proBDNF, and another detection antibody against the N-terminal of mBDNF. This allowed us to measure both mBDNF and proBDNF levels simultaneously in the same sample. Unfortunately, our highly sensitive ELISA assay (down to 2 pg/ml) could not detect any trace amount of mBDNF nor proBDNF with the 10 ul of mouse blood. Consistent with this, several previous reports indicated that, unlike human and rat blood, mouse blood does not have BDNF (Klein et al., 2011; Radka, Holst, Fritsche, & Altar, 1996).

We therefore turned to examine the levels of mBDNF and proBDNF in the plasma of human ASD patients and healthy controls. We obtained 1 ml of blood from human volunteers (controls), 100 times the amount from mice. In the initial, pilot experiments, 9 ASD patients and 10 healthy volunteers were enrolled, with well-documented consent forms and the approval of the Institutional Ethics Board. The age of participants ranged from 3∼13 years: 7 boys and 3 girls in the ASD group; 1 boy and 10 girls in the control group (Table 1). Among those 10 ASD patients, all had delayed development of speech, and most showed repetitive behaviors, a common feature in ASD diagnosis. We also noticed that the intellectual abilities of at least 8 out of 10 ASD children were also impaired, suggesting that these ASD patients likely belonged to the traditional ASD, but not some special groups of ASD with normal intellectual ability, such as Asperger’s syndrome.

**Table 1.** The general information of ASD patients and controls. Subject ID, gender, birthday, recruit data, and age of all ASD patients and controls were displayed.

| Subject ID | Gender | Birthday | RecruitDate | Age (year) |
| --- | --- | --- | --- | --- |
| ASD |  |  |  |  |
| HN0539.p1 | M | 2015/11/27 | 2018/12/9 | 3.04 |
| HN0543.p1 | M | 2013/10/6 | 2018/12/9 | 5.18 |
| HN0544.p1 | M | 2015/4/27 | 2018/12/9 | 3.62 |
| HN0555.p1 | F | 2016/2/3 | 2018/12/16 | 2.87 |
| HN0557.p1 | F | 2009/8/21 | 2018/12/16 | 9.33 |
| HN0560.p1 | F | 2013/11/21 | 2019/1/4 | 5.12 |
| HN0562.p1 | M | 2016/7/8 | 2019/3/20 | 2.70 |
| HN0612.p1 | M | 2017/6/30 | 2019/7/21 | 2.06 |
| HN0620.p1 | M | 2016/11/19 | 2019/7/21 | 2.67 |
| HN0620.p2 | M | 2016/11/19 | 2019/7/21 | 2.67 |
| Control |  |  |  |  |
| HN0539.s1 | F | 2012/1/1 | 2018/12/9 | 6.94 |
| HN0543.s1 | F | 2016/8/27 | 2018/12/9 | 2.28 |
| HN0544.s1 | F | 2007/8/23 | 2018/12/16 | 11.32 |
| HN0555.s1 | F | 2013/3/13 | 2018/12/16 | 5.76 |
| HN0555.s2 | F | 2016/2/3 | 2018/12/16 | 2.87 |
| HN0557.s1 | F | 2005/6/17 | 2018/12/16 | 13.51 |
| HN0557.s2 | F | 2009/8/21 | 2018/12/16 | 9.33 |
| HN0560.s1 | F | 2006/9/2 | 2019/1/4 | 12.35 |
| HN0562.s1 | F | 2014/12/9 | 2019/3/20 | 4.28 |
| HN0612.s1 | M | 2015/6/26 | 2019/7/21 | 4.07 |
| HN0620.s1 | F | 2013/11/11 | 2019/7/21 | 5.69 |

Next, peripheral venous blood, from patients and controls. was collected in EDTA-coated tubes and centrifuged within 5 min to obtain plasma (see Methods). Using the sandwich ELISA specific for proBDNF and mBDNF, respectively, we found a selective increase in proBDNF, but not mBDNF, in the ASD patients (Figure 7, and Table 6). The concentration of proBDNF was 1316.17 pg/ml, whereas that of mBDNF was 946.27 pg/ml in ASD patients. In healthy controls, the concentration of proBDNF was 688.39 pg/ml, nearly half of that in the ASD group. However, the concentration of mBDNF is 713.12 pg/ml, which was comparable to that in the ASD group. Thus, the increase in plasma proBDNF, but not mBDNF, is associated with ASD.

**Figure 7.**
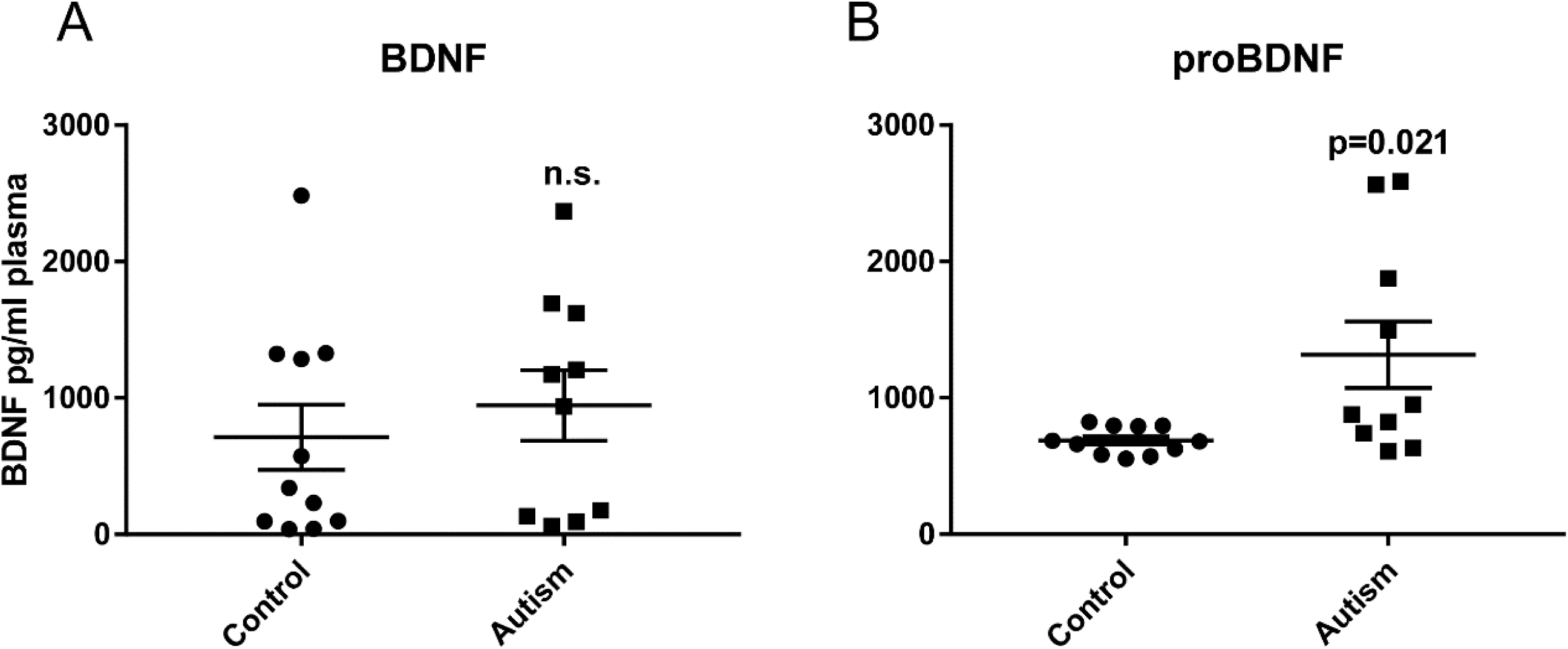
proBDNF and BDNF levels in human plasma. (**A, B**) ELISA analysis of mBDNF (**A**) and proBDNF (**B**) in human plasma of ASD (n=11) and control (n=10) children. Blood sampling and the separation of plasma was described in Material and Methods. A sandwich ELISA was employed to detect both mBDNF and proBDNF levels simultaneously. Data are shown as mean ± S.E.M.

Furthermore, we observed, evaluated, and documented all related symptoms of these ASD patients (Table 2∼5). 8 out of 10 patients were firmly diagnosed with ASD, while in the remaining two, autistic features were clearly found, but these features did not match the criteria of ASD diagnosis (Table 2). All patients exhibited conventional autistic features, including the delay of speech development (Table 2), intellectual disabilities (Table 2), repetitive behavior (Table 4), and ADHD (Table 4). However, in some of these ASD patients, we also observed unconventional symptoms, including delays in motor development (Table 2), sleep disturbances (Table 3), anxiety (Table 4), self-injury (Table 4), obsessive behavior (Table 4), and aggressive behavior (Table 4). Interestingly, we found that the proBDNF data of ASD patients was associated with a few of these unconventional symptoms. Several ASD children with aggression, eye abnormalities, sleep disorders, and self-injurious behavior tended to express lower proBDNF levels (patient HN0544.p1, HN0555.p1, HN0557.p1, and HN0560.p1). This observation may imply that proBDNF is more likely a biomarker only for ASD with common features, rather than all the ASD subtypes. Further studies with a larger patient number are necessary to validate this point.

**Table 2.** Neurodevelopment problems in the ASD patients. Neurodevelopment symptoms of the ASD patients, including developmental delay of speech and motor ability, ASD or autisitc features, and intellectual disability were summarized. In this and all the following tables, ‘+’ represents a positive symptom in an ASD patient, ‘-’ represents a negative one, and the blank means the imformation is not reported. Please note ‘±’ means the patient exhibit autistic features but not severe enough to be disagoised as the ASD.

| Subject ID | developmental delay (speech) | developmental delay (motor) | ASD/autistic features | Intellectual disability |
| --- | --- | --- | --- | --- |
| HN0539.p1 | + |  | ± | + |
| HN0543.p1 | + | + | + | + |
| HN0544.p1 | + | — | + | + |
| HN0555.p1 | + | + | + | + |
| HN0557.p1 | + |  | + |  |
| HN0560.p1 | + |  | + |  |
| HN0562.p1 | + |  | + | + |
| HN0612.p1 | + | — | ± | + |
| HN0620.p1 | + | — | + | — |
| HN0620.p2 | + | — | + | + |

**Table 3.** Neurological problems in the ASD patients. Neurological tests for epilepsy, EEG and MRI abnormities, macrocephaly and sleep disorders in the ASD patients. Please note ‘±’ in the column of epilepsy means seizure was found in that patients, but not severe enough for epilepsy diagnosis.

| Subject ID | Epilepsy /Seizure | EEG abnormalities | MRI Brain abnormalities | Macrocephaly | Sleep disturbances |
| --- | --- | --- | --- | --- | --- |
| HN0539.p1 |  |  |  | — | — |
| HN0543.p1 | — | — | — | — | — |
| HN0544.p1 | — | — | — | — | — |
| HN0555.p1 | — | — | — | — | — |
| HN0557.p1 |  |  |  | — | + |
| HN0560.p1 |  |  |  | — | + |
| HN0562.p1 |  |  |  | — | — |
| HN0612.p1 | — | — | — | — | — |
| HN0620.p1 | — |  |  | — | — |
| HN0620.p2 | — |  |  | — | — |

**Table 4.** Behavioral problems in the ASD patients. Behavior problems, including, the repetitive behavior, ADHD, anxiety, obsessive behavior, self-injury, and aggressive behavior, were displayed in the ASD patients.

| Subject ID | Repetitive behavior | ADHD | Anxiety | Obsessive behavior | Self-injurious behavior | Aggressive behavior |
| --- | --- | --- | --- | --- | --- | --- |
| HN0539.p1 | + |  |  | — | — | — |
| HN0543.p1 | + | + | — | + | — | — |
| HN0544.p1 | + | + | + | — | — | — |
| HN0555.p1 | + | + | — | — | — | — |
| HN0557.p1 | + | + | + |  | + | — |
| HN0560.p1 | + | + |  |  | — | + |
| HN0562.p1 | + | + |  | + | + | — |
| HN0612.p1 |  | — | — | — | — | — |
| HN0620.p1 | + | — | — | — | — | — |
| HN0620.p2 | + | — | — | — | — | — |

**Table 5.** No systemic problems observed in the ASD patients. Systemic features were documented for these ASD patients. No systemic problems, such as eye abnormalities, recurrent infections, hypotonia, hand deformity, and short stature were observed in all the ASD patients.

| Subject ID | Eye abnormalities | Recurrent infections | Hypotonia | Hand deformity | Short stature |
| --- | --- | --- | --- | --- | --- |
| HN0539.p1 | — |  |  | — | — |
| HN0543.p1 | — | — | — | — | — |
| HN0544.p1 | — | — | — | — | — |
| HN0555.p1 | — | — | — | — | — |
| HN0557.p1 |  |  |  | — | — |
| HN0560.p1 |  |  | — | — | — |
| HN0562.p1 | — |  | — | — | — |
| HN0612.p1 | — | — | — | — | — |
| HN0620.p1 | — | — | — | — | — |
| HN0620.p2 | — | — | — | — | — |

## Discussion

Most of the previous models for ASD were based on SNPs or gene mutations found in ASD patients. However, emerging studies have shown that most of ASD patients are not familial but sporadic, and genetic association studies have identified numerous risk variations but few causal genes (De La Torre-Ubieta et al., 2016). Interestingly, nearly all well-founded ASD related genes, such as *fmr1, mecp2, scna1*, and *cacnal1*, are associated with synaptic function (Amir et al., 1999; Bassell & Warren, 2008; De Rubeis et al., 2014; Mahoney et al., 2009; Oakley et al., 2011; Splawski et al., 2004). Furthermore, most ASD animal models exhibit reductions in synaptic density, synaptic protein synthesis, or synaptic plasticity. In this study, we took an unconventional approach, focusing not on genetic mutations but on the processing of BDNF, a neurotrophic protein known to play a critical role in synaptic function. We have developed an ASD-like mouse model in which synaptic function is decreased, through elevating the proBDNF level but reducing mBDNF level. To the best of our knowledge, this is the first ASD mouse model not based on genetic mutations, but on protein processing.

Due to the lack of tools to non-invasively detect cellular changes in human brain, we do not know the neuronal phenotypes in the brains of ASD patients. However, studies using postmortem ASD brain tissues have revealed increased spine density in many brain regions, such as the frontal, parietal and temporal lobes (Hutsler & Zhang, 2010; Tang et al., 2014). Efforts have been made to study neuronal phenotypes using ASD animal models. Compared with some of previous models, our BDNF^met/leu^ mice exhibit more comprehensive neuronal phenotypes relevant to ASD. Specifically, the BDNF^met/leu^ mice showed reduced brain volume and dendritic complexity, increased thin spines but decreased mushroom spines, altered synaptic proteins, decreased basal synaptic transmission, and impairments in both LTP and LTD. While previously published ASD models display some synaptic deficits, there is no model that express all of these phenotypes (De La Torre-Ubieta et al., 2016). The similarity in the synaptic phenotypes between our BDNF^met/leu^ mice and previous ASD models suggests that processing of BDNF protein, rather than genetic alteration of *Bdnf* gene, may be an important factor contributing to ASD etiology. However, animal and human studies have so far not implicated BDNF in ASD. Thus, our study provides the first link between ASD and BDNF. Given that BDNF is a key regulator of synaptic development and function (Lu, Pang, & Woo, 2005a), the present work also opens up a new direction for ASD research.

Another important finding is that the BDNF^met/leu^ mice exhibit comprehensive behavior deficits resembling humans with ASD. We showed the BDNF^met/leu^ mice have impaired social interaction and repetitive behaviors, two key components in the diagnostic criteria for ASD as defined by DSM-5. Compared with all previously established ASD models, the repetitive behaviors of the BDNF^met/leu^ mice are much stronger and more like the symptoms of human ASD patients. A unique aspect of the BDNF^met/leu^ mice, not seen in other ASD models, is the strong and spontaneous ‘star-gazing’ stereotype (Movie #1), in the absence of any external stimuli (‘self-stimulating’). In many children with ASD, especially for those diagnosed with Tourette’s syndrome, the ‘self-stimulating’ behaviors are extremely frequent and apparent, such as flapping their arms or wiggling their toes repeatedly(Kurlan, 2010; Leckman, 2002), which are categorically similar to the ‘star-gazing’ stereotype in the BDNF^met/leu^ mice. It should be noted that the BDNF^met/leu^ mice do not show over-grooming, a form of repetitive behavior observed in some ASD models but not in human ASD. Bead-burying, a repetitive behavior seen in some ASD models but not in ASD children, is also absent in our BDNF^met/leu^ mice. Thus, the BDNF^met/leu^ mice could be a different and perhaps better ASD model, with regards to ‘self-stimulating’ behaviors. Regardless, the robust behavioral phenotype of the BDNF^met/leu^ mice provides a unique opportunity for mechanistic studies and drug testing.

Finally, our preliminary data indicates that the plasma level of proBDNF could be used as a potential ASD biomarker that may help in monitoring disease progression and drug efficacy. Specifically, in the pilot experiment with 9 ASD patients and 10 healthy volunteers, we found that an increase in plasma proBDNF level, but not mBDNF level, is associated with ASD. However, a biomarker for clinical use requires both good sensitivity and specificity (Brower, 2011). Our data (Figure 7 and Table 6) indicates that the concentration of proBDNF in the ASD group is twice of that in healthy controls, suggesting that the sensitivity of the current ELISA might be in the right range. Technologies exist to further improve the sensitivity greatly. Given the small sample size, we cannot claim the specific association of elevated proBDNF with ASD. Further studies with much larger sample sizes are necessary to validate the specificity of this potential biomarker and to investigate whether it is associated with a distinct ASD subtype. Previous studies have suggested that the plasma level of BDNF could reflect BDNF concentration in the brain (Klein et al., 2011). It is unclear whether the plasma level of proBDNF also correlates with proBDNF concentration in the brain. The source and the functional implication of plasma proBDNF should also be investigated in the future.

**Table 6.** The mBDNF and proBDNF level of plasma in the ASD patients. The mBDNF and proBDNF level of plasma in each of ASD patients were displayed.

| Subject ID | mBDNF level (pg/ml) | proBDNF level (pg/ml) |
| --- | --- | --- |
| HN0539.p1 | 1172.76 | 1877.297 |
| HN0543.p1 | 132.7449 | 1495.534 |
| HN0544.p1 | 1693.265 | 608.195 |
| HN0555.p1 | 175.8991 | 824.8709 |
| HN0557.p1 | 92.91032 | 633.9897 |
| HN0560.p1 | 60.37873 | 740.608 |
| HN0562.p1 | 1621.231 | 878.18 |
| HN0612.p1 | 938.3997 | 950.4053 |
| HN0620.p1 | 1206.951 | 2589.232 |
| HN0620.p2 | 2368.13 | 2563.437 |

Although most of the phenotypes found in the BDNF^met/leu^ mice coincide with the symptoms of ASD patients or neuronal features of the other ASD models, the brain volume changes may not be the same as those found in human studies. In contrast to the decrease of brain volume in the BDNF^met/leu^ mice, previous studies with large cohorts have shown brain overgrowth in ASD children (De La Torre-Ubieta et al., 2016). However, no such brain overgrowth was reliably detected in the adult ASD patients. In addition, neuroimaging analysis of ASD brains also showed decreased volume of the corpus callosum, cerebellum and brainstem (Penn, 2006). These clinical observations suggest that brain volume may not be used as a general pathological criterion for ASD. Interestingly, mutations of the X-linked genes encoding neuroligins, which are known for their association with ASD, have been shown to lead to decreased brain volume (Jamain et al., 2003). Moreover, both neuroligins-3 and −4 knock-out mice, which could display all ASD-like phenotypes, have reduced brain volume (Baudouin et al., 2012; Jamain et al., 2008). Thus, our BDNF^met/leu^ mice could be an ASD-like model similar to fragile X syndrome.

Another phenotype which shows a potential inconsistency with a previous finding is the reduced LTD of the BDNF^met/leu^ mice. A previous study found an increase, not decrease, in hippocampal LTD in the heterozygotes of cleavage-resistant *probndfHA* knock-in mice (Yang et al., 2014). The design of *probndfHA* knockin line was such that the last two arginines on the proBDNF cleavage site was replaced by two alanines (RR→AA), and therefore, the cleavage of proBDNF was completely blocked. Since the homozygous *probndfHA* die at birth, only the heterozygous mice were used for the LTD experiment. The heterozygous *probndfHA* knock-in line contains one allele with uncleavable proBDNF and another WT allele. Thus, the proBDNF/mBDNF ratio should be approximately 1:1. The increased LTD seen in *probndfHA* heterozygotes is consistent with the observation that application of exogenous proBDNF to WT hippocampal slices facilitates LTD (Woo et al, 2015). In contrast, our BDNF^met/leu^ mice are homozygous and the mBDNF level in this line was decreased by nearly 90% of that in the WT mice. The extremely low level of mBDNF in the BDNF^met/leu^ mice may not maintain normal function and morphology of hippocampal spines and thus results in a decrease rather than an increase in LTD. This may explain the difference in LTD phenotype between our BDNF^met/leu^ mice and the *probndfHA* heterozygous mice.

In summary, we have developed a new ASD model, not based on genetic mutations, but on protein processing, and provided a link between ASD and BDNF, a key factor for synaptic regulation. With its robust phenotypes, the BDNF^met/leu^ mouse line may serve as a new ASD model with more comprehensive behavioral deficits resembling human autism patients. Finally, our preliminary study using human blood samples raises the possibility of using plasma proBDNF level as an ASD biomarker.

## Materials and Methods

### Animals

All animal experiments were performed in strict accordance with protocols that were approved by the Institutional Animal Care and Use Committee of AIST. All efforts were made to minimize animal suffering during the experiments. Animals

All animal experiments were performed in strict accordance with protocols that were approved by the Institutional Animal Care and Use Committee of AIST (number.2019-084, approval date 6 June 2015). All efforts were made to minimize animal suffering during the experiments. Mice were housed with access to food and water ad libitum at constant room temperature (24 ± 2°C) and were exposed to a 12 h light/dark cycle (lights on between 7 a.m. and 7 p.m.). All mice were housed in groups of 3–4 per cage.

### Generation of proBDNF knock-in mouse line

Generation of proBDNF knock-in mouse line was described in the report of Kojima et al. (2020) (Kojima et al., 2020). Briefly, the proBDNF knock-in mouse line was generated by replacing the endogenous Bdnf allele with a cDNA encoding cleavage-resistant form of BDNF that contained two amino acid substitutions proximal to the cleavage site of proBDNF (Figure 1A, RVRR to MVLR). A 6.0 kb long arm fragment (NCBI accession number: AY057907; 49015–54460) and a 3.4 kb short arm fragment (55207–58754) flanking the 5’ and 3’ ends of mouse Bdnf exon 5, respectively, were amplified from 129SV mouse genomic DNA using PCR. The long and short arm fragments were introduced into the ClaI-NotI and PacI-AscI sites of the pMulti-ND 1.0 vector, respectively. A DNA fragment encoding mutant proBDNF was then inserted into the ClaI-NotI site of the targeting vector. A pGK-thymidine kinase gene was used as a negative selectable marker (Figure 1A, Vector). Linearized targeting vector was electroporated into embryonic stem (ES) cells of the D3 line (strain 129 SV). DNA derived from G418-resistant ES clones was screened using PCR and positive ES clones were injected into blastocysts obtained from C57BL/6 mice. The injected blastocysts were then introduced into the uteri of pseudo-pregnant females. Chimeric mice were mated with C57BL/6 mice to produce heterozygotes, and these mice were subsequently crossed with mice expressing Cre recombinase in germ cells to excise the neo cassette. A genetic backcross to the 129/SvEv background were performed over 10 generations before the animals were used in behavioral analyses and all other analyses.

### Genotyping

Genotyping of the proBDNF knock-in mouse line was performed according to our recent report (Kojima et al., 2020). using the following primers: 5’-TGCACCACCAACTGCTTAG-3’ and 5’-GGATGCAGGGATGATGTTC-3’. PCR analyses using these primers generated 550 bp and 320 bp DNA fragments from wild-type and mutant alleles, respectively. Genotyping of the p75NTR knock-out mice (Ngfrtm1Jae) from the Jackson Laboratory (Bar Harbor, ME) was performed using the following primers: 5’-GCTCAGGACTCGTGTTCTCC-3’, 5’-CCAAAGAAGGAATTGGTGGA-3’, and 5’-TGGATGTGGAATGTGTGCGAG-3’. PCR analyses using these primers generated 386 bp and 193 bp DNA fragments from wild-type and mutant alleles, respectively. Genotyping of the BDNF knock-out mice (kindly provided by Prof. Nakamura, Tokyo University of Agriculture and Technology, Tokyo) was performed using the following primers: 5’-ATGAAAGAAGTAAACGTCCAC-3’, 5’-CCAGCAGAAAGAGTAGAGGAG-3’, and 5’-GGGAACTTCCTGACTAGGGG-3’. PCR analyses using these primers generated 275 bp and 340 bp DNA fragments from wild-type and mutant alleles, respectively. Wild-type mice with the background of 129S6/SvEv and C57BL6/J were obtained from Taconic Biosciences (Hudson, NY) and Crea Japan (Shizuoka, Japan), respectively.

### Golgi impregnation of granule neurons in the hippocampal dentate gyrus

Golgi impregnation of mouse brains was performed using the FD Rapid GolgiStain Kit. Dentate gyrus (DG) neurons were examined in the dorsal hippocampus. For Sholl analyses of dendritic arborization (Z. Y. Chen et al., 2006), Golgi-impregnated DG granule cells that met the following criteria were used: (i) isolated cell body with a clear relationship between the primary dendrite and the soma, (ii) presence of untruncated dendrites, (iii) consistent and dark impregnation along the extent of all of the dendrites, and (iv) relative isolation from neighboring impregnated cells that could interfere with the analysis. For quantitative analyses, 50 neurons from the hippocampal DG area were selected. The cells were traced under 40× magnification using Neurolucida software (MicroBrightField Inc., Colchester, VT) and the morphological traits were analyzed using the NeuroExplorer analysis package. The data were processed and statistical analyses were performed using Graph Pad Prism 4.0 (Graph Pad Software Inc., San Diego, CA).

### Cavalieri analysis of Nissl-stained sections

Cavalieri analysis of Nissl-stained sections were used for hippocampal volume estimation. Steroinvestigator software was applied to measure the entire volume of the hippocampus at 4X objective magnification. The external capsule, alveus of hippocampus, and white matter were used as boundary landmarks. All sections throughout each hippocampus were traced and reconstructed. The Cavalieri estimator function was used to calculate the volume of each hippocampus. Following total hippocampal measurements, the cellular layer of each subregion of the hippocampus (DG, CA1, CA2/3) was traced separately and analyzed in the same manner.

### Immunohistochemistry

Adult mice were anesthetized and transcardially perfused with phosphate buffered saline (PBS) followed by 4% paraformaldehyde in PBS. The brains were postfixed overnight at 4°C, cryoprotected, and then cut into 30 µm sections. The sections were washed three times in PBS for 10 min, permeabilized with 0.2% Triton X-100 in PBS, washed a further three times in PBS for 10 min, incubated in 3% bovine serum albumin (BSA) in PBS for 2 h, and then incubated at 4°C for 48 h with anti-proBDNF rabbit antiserum(H. Koshimizu et al., 2009) diluted 1:500 in PBS containing 3% BSA. Subsequently, the sections were washed three times in PBS for 5 min and then incubated with Alexa555 goat anti rabbit IgG (1:1000, Life Technologies, Carlsbad, CA) and 4’,6’-diamidino-2-phenylindole dihydrochloride (1 µg/ml, Life Technologies) for 2 hr. Fluorescent images were obtained using a Nikon C1 confocal or Nikon Ti E inverted microscope (Nikon, Japan) and a 20× Plan Apo, NA 0.75 numerical aperture objective lens (Nikon). Immunocytochemical staining using a rat monoclonal anti-mouse CD31 antibody (1:100; clone MEC 13.3; BD Pharmingen, San Jose, CA) was used to detect endothelial cells in the mouse heart. Purified anti-phospho-TrkB (pTrkB) antiserum, which was generously provided by Dr. Chao(Arevalo et al., 2006), was used to detect the activated TrkB receptor. Mouse heart sections were treated with 0.1% hydrogen peroxide and then incubated with the primary antibody. Signal amplification was performed using the avidin:biotinylated horseradish peroxidase complex method (Vectastain ABC; Vector Labs, Burlingame, CA). An anti-serotonin (5-HT) antibody (1:15,000; Incstar, Stillwater, MN) was used for immunocytochemical analyses of 5-HT in brain tissues and the fiber density was quantified as described by Lyons et al(Lyons et al., 1999).

### In situ hybridization

Brains from 8-week-old mice were fixed by transcardial perfusion with PBS followed by 4% paraformaldehyde in PBS at 4°C. The brains were immersed in the fixative overnight at 4°C, treated with 20% sucrose in PBS, embedded in Tissue-Tek Optimal Cutting Temperature Compound (Sakura Finetek, Tokyo, Japan), and then frozen on a block of dry-ice. In situ hybridization was performed on 20 µm coronal sections. The hybridization and post-hybridization wash steps were performed at 65°C to avoid nonspecific binding. Stained sections were photographed under the bright field system of a BioRevo microscope (Keyence, Osaka, Japan).

### Immunoblotting

Immunoblotting was performed according to our previous report(Suzuki et al., 2007). Briefly, hippocampal tissues were homogenized in cold lysis buffer comprising 50 mM Tris-HCl (pH 7.4), 1 mM EDTA, 150 mM NaCl, 10 mM NaF, 1 mM Na3VO4, 1% Triton X-100, 10 mM Na2P2O7, 100 µM phenylarsine oxide, and 1% protease inhibitor cocktail (Complete Mini, Roche Diagnostics, Hertforshire, UK). The lysed tissues were incubated at 4°C for 20 min and then centrifuged at 15,000 rpm for 15 min. The protein levels in the supernatants were determined using the BCA Protein Assay Kit (Pierce/Thermo Scientific, Rockford, IL). To detect BDNF, the lysates were boiled for 5 min at 100°C, separated on SDS-polyacrylamide gels, and then transferred to polyvinylidene fluoride membranes (Immobilon P; Millipore, Billerica, MA). The membranes were blocked in Tris-buffered saline containing 0.2% Tween-20 (TBST) and 5% BSA or Block Ace (Dainippon Pharmaceutical, Osaka, Japan), incubated with a rabbit polyclonal anti-BDNF antibody (N-20, Santa Cruz Biotechnology, San Diego, CA) in TBST containing 0.5% BSA or 1% Block Ace at room temperature for 90 min, and then washed three times with TBST. Subsequently, the membranes were incubated at room temperature for 30 min with peroxidase-conjugated secondary antibodies in TBST containing 5% BSA and washed three times with TBST. The signal was detected using ImmunoStar Reagents (Wako, Tokyo, Japan) or SuperSignal WestFemto Maximum Sensitivity Substrate (Pierce). The exposure time was adjusted such that the intensity of the bands was within the linear range. To quantify the amount of proBDNF relative to the amount of total BDNF, the blots were scanned and the images were converted to TIFF files and quantified using the NIH ImageJ software (version 1.37v). After subtracting the background in the same lane, the signals for proBDNF and mature BDNF were summed to obtain the total amount of BDNF.

To detect other molecules, the lysates were separated on SDS-polyacrylamide gels and transferred to polyvinylidene fluoride membranes (Immobilon P membrane, Millipore). The membranes were blocked at room temperature for 30 min in TBST containing 5% skimmed milk (Nacalai Tesque, Kyoto, Japan), and then incubated with the primary antibodies in TBST containing 3% BSA at room temperature overnight. The following primary antibodies were used: anti-p75NTR (1:1000; clone D4B3; Cell Signaling Technology, Danvers, MA), anti-SorCS2 (1:500; R&D Systems, Minneapolis, MN), anti-SorCS3 (1:500; R&D Systems), anti-PSD-95 (1:1000; clone D27E11; Cell Signaling Technology), anti-SynGAP (1:1000; clone D88G1; Cell Signaling Technology), anti-GluN2B (1:1000; Alomone Labs, Jerusalem, Israel), anti-synapsin I (1:1000; Millipore), and anti-β-actin (1:10000; clone AC-15; Sigma, St. Louis, MO). After washing, the membranes were incubated with TBST containing horseradish peroxidase-conjugated secondary antibody (1:1000; GE Healthcare, Pittsburgh, PA) for 1 h at room temperature. The blots were developed using Luminata Forte Western HRP substrate (Millipore) and images of the immunoblots were captured using an ImageQuant LAS500 (GE Healthcare) imaging system. Quantitative analyses of the band intensities were performed using ImageQuant software (Image Analysis Software v8.1, GE Healthcare). Hippocampal tissues were isolated from 9-week-old mice and rapidly homogenized in lysis buffer comprising 50 mM Tris-HCl (pH 7.4), 1 mM EDTA, 150 mM NaCl, 10 mM NaF, 1 mM Na3VO4, 1% Triton X-100, 10 mM Na2P2O7, 100 µM phenylarsine oxide, 1% protease inhibitor cocktail, and 1% protein phosphatase inhibitor cocktail (Sigma). Lysed tissues were incubated at 4°C for 20 min and centrifuged at 15,000 rpm for 15 min. The protein levels in the supernatants were determined using the BCA Protein Assay kit (Pierce). Protein G Sepharose (50 µl, GE Healthcare) was added to the hippocampal lysates and the mixture was rotated at 4°C for 60 min. After removing the Protein G Sepharose, 1 µg of a rabbit monoclonal anti-p75NTR antibody (clone D4B3; Cell Signaling Technology) or mouse monoclonal anti-PSD-95 antibody (clone 7E3-1B8; Thermo Scientific, Pittsburgh, PA) was added to the supernatant (containing 2.5 mg of total protein). The lysates were incubated with the beads (50 µl) at 4°C overnight. The immune complexes were then pelleted and immunoblotted with mouse monoclonal anti-PSD-95 (1:1000, Thermo Scientific), rabbit monoclonal anti-PSD-95 (1:1000; clone D27E11; Cell Signaling Technology), or rabbit monoclonal anti-SynGAP (1:1000; clone D88G1; Cell Signaling Technology).

### Golgi staining and quantitative analyses of dendritic spine morphology

Mice were anesthetized with isoflurane and the brains were removed and washed in ice-cold PBS (pH 7.4). Rapid Golgi staining was performed using the FD Rapid GolgiStain Kit (FD NeuroTechnologies, Ellicott City, MD), according to the manufacturer’s instructions. Briefly, whole brains were silver impregnated for 2 weeks, cryoprotected for 1 week, and then sectioned (80 m) on a sliding microtome (REM-700, Yamato Kohki. Industrial Co., Ltd., Saitama, Japan). The sections were developed, clarified, and then coverslipped with resinous medium. During staining, image acquisition, and image analysis, the operators were blinded to the genotype of each animal. Images of secondary dendritic segments of hippocampal pyramidal neurons in the stratum radiatum in the CA1 region were obtained using an AxioObserver Z1 microscope (Zeiss, Oberkochen, Germany) with a 63×/0.75 numerical aperture objective lens (LD PlanNeo; Zeiss) for high magnification imaging or a 20×/0.8 numerical aperture objective lens (PlanApo; Zeiss) for low magnification imaging, and a cooled CCD camera (CoolSnap HQ2; Photometrics, Tucson, AZ). For morphological analyses of dendritic spines, a z-series with 0.5 µm intervals was created to visualize the entire extent of each dendritic process and to capture spine details in all focal planes. The density, maximum length, and maximum width of the dendritic protrusions was quantified using the Region Measurements tool of MetaMorph Software (Universal Imaging Corporation, West Chester, PA, USA) (10). The form factor of each protrusion, defined as the ratio of its maximum length to its maximum width, was calculated(Takahashi et al., 2003). The age, genotype, and number of mice used for morphological analyses are described in the figure legends. The morphologies of dendritic protrusions were quantified using 33–36 segments of the secondary dendrites. These segments represented a total dendritic length of 1329–1683 µm per group of mice.

### Electrophysiology

Field excitatory synaptic potentials were recorded as follows. Transverse slice was placed on custom-made recording chamber according to a previous report (Tominaga, Tominaga, & Ichikawa, 2002) with some modifications. Briefly, both sides of slice were perfused with carbogenize aCSF containing (in mM) 118 NaCl, 3 KCl, 2.5 CaCl2, 1.2 MgCl2, 11 D-glucose, 10 HEPES, and 25 NaHCO3, at 2-3ml/min at 28 C. Field EPSPs were elicited by 10-100 μA constant currents pulse (100 microsecond duration) with Pt bipolar electrode (FHC) onto the Schaffer-collateral pathway and recored with glass electrode filled with aCSF (1-2 M ohm). EPSPs were amplified by multiclamp700A (Axon instruments, Sunnydale CA, USA) and digitized at 10Hz (National Instruments, Austin, Texas, USA) controlled by WinLTP programs (Anderson, Fitzjohn, & Collingridge, 2012).

To set stimulus intensities, maximum responses were determined by increasing stimulus intensity stepwise to saturating responses. For paired-pulse experiment and long-term potentiation experiments, stimulus intensity which elite about 33% of maximum response were employed. Input-output relationship were studied by plotting fiber volley response amplitude (mV) against fEPSP slope (mV/ms). Paired-pulse responses were also recorded with various inter-pulse intervals from 500ms, 100ms, 50ms, 20ms and paired-pulse ration (2nd pulse response to first response) were calculated. LTP was induced by four theta burst stimulation, which consist of 4 burst at 0.05Hz of theta burst stimuli (4 bursts at 10Hz of 5 pulse at 100Hz). On the other hands, LTD was induced by low frequency simulation (1Hz 900 pulse). In case of LTD experiments, stimulus intensity which elicited 50 % of maximum response was employed.

### Blood sampling and corticosterone measurements

Corticosterone measurements were performed as described previously(Schmidt et al., 2007). To determine basal corticosterone levels, blood samples were collected between 10 and 12 a.m. The mice were moved to an adjacent room and placed individually in a transparent restraint tube. Blood samples (approximately 100 µl) were collected into plastic tubes from the tail vein without anesthesia, as described previously(Fluttert, Dalm, & Oitzl, 2000). In all cases, the time between the first disturbance of the animals and the sampling was less than 1 min. Following blood collection, the mice were released and returned to their cages. Blood samples were separated by centrifugation at 4°C for 15 min and the plasma was stored in the freezer until use. To measure the change in blood corticosterone levels in response to acute stress, sampling was performed 60 min before and after restraint stress was initiated. Restraining was performed in plastic cylinders identical to those used for the prenatal stress procedure. The AssayMax Corticosterone ELISA Kit (Assaypro, St. Charles, MO) was used to measure corticosterone levels in the blood samples, according to the manufacturer’s instructions.

### Reverse transcription and quantitative PCR

Hippocampal tissues from adult (8–15-week-old) mice (single mutants: BDNF^+/+^, n = 9; and BDNF^met/leu^, n = 8; and double mutants: BDNF^+/+^; p75^NGFR+/+^, n = 6; BDNF^+/+^; p75^NGFR+/-^, n = 7, BDNF^met/leu^; p75^NGFR+/+^, n = 7; and BDNF^met/leu^; p75^NGFR+/-^, n = 6) were dissected and stored in RNAlater (Applied Biosystems, Foster City, CA) at −30°C until use. Total RNA was extracted using ISOGEN reagent (Nippon Gene, Tokyo, Japan), treated with RNase-free DNase I (Qiagen, Valencia, CA) to remove contaminating genomic DNA, and then purified using the RNeasy Mini Kit (Qiagen), according to the manufacturer’s instructions. Total RNA (0.5 μg) was reverse-transcribed into cDNA using SuperScript III (Invitrogen, Carlsbad, CA) and an oligo (dT)20 primer or BDNFrt-3 primer (for *Bdnf* antisense-specific PCR) in a total volume of 5 µl. Quantitative real-time PCR was performed using the 7300 Real-Time PCR system (Applied Biosystems). Duplicate standards were included in each plate and each sample was measured in triplicate. PCR was performed in a total volume of 15 µl containing 1.5 µl of diluted cDNA (1/80 dilution), specific forward and reverse primers (50 nM), and Power SYBR Green PCR Master Mix (Applied Biosystems). The PCR conditions for all genes were as follows: 50°C for 2 min, 95°C for 10 min, and then 40 cycles of 95°C for 15 s and 60°C for 1 min. All mRNA levels were normalized to those of Gapdh to adjust for small differences in the amount of input RNA. The primer sequences were as follows: BDrt-1 (for BDNF mRNA), 5’-TGTGTGACAAGTATTAGCGAGTGG-3’; BDrt-2 (for BDNF mRNA), 5’-ATGGGATTACACTTGGTCTCGT-3’; BDrt-3 (for BDNF antisense RNA), 5’-ACTCTGGAGAGCGTGAATG-3’; BDrt-4 (for BDNF antisense RNA), 5’-GGCTCCAAAGGCACTTGAC-3’; NGF-F, 5’-GATGGCATGCTGGACCCAAG-3’; NGF-R, 5’-CAACATGGACATTACGCTATGCAC-3’; NT3-F, 5’-GGATGATGACAAACACTGG-3’; NT3-R, 5’-ACAAGGCACACACACAGGAA-3’; GAPDH-F, 5’-TGCACCACCAACTGCTTAGC-3’; and GAPDH-R, 5’-GGCATGGACTGTGGTCATGAG-3’. The expression levels of genes in mutant tissues were normalized to those of the corresponding genes in wild-type tissues.

### Tail-suspension test (TST) and antidepressant treatment

For tail-suspension test (TST) (Figure 6), Male mice at 8-12-week-old with the indicated genotype were used. TST was performed on an automated tail suspension device (Muromachi Kikai, Tokyo, Japan). Animals were suspended from a strain gauge for 6 minutes. The settings for the equipment were time constant= 0.25, gain=4, threshold 1=3, and resolution=200 ms. Time spent immobile was recorded in seconds. To test the effect of antidepressant treatment on depressive-behavior, animals were administered saline or fluoxetine 30 min before the tail suspension test (10 mg/kg i.p.).

### Human plasma collection

All human plasma collection protocols were approved by the Ethics Committee at Xiangya Hospital Central South University. 11 Autism children and their 10 normal siblings as control were recruited for blood sampling. Peripheral venous blood from patients and controls was collected in EDTA-coated tubes and centrifuged at 1000g, 10 minutes to obtain the raw plasma. Then, the raw plasma was centrifuged at 1000g, 10 minutes again to eliminate residual blood cells. All plasma samples were stored at −80°C before the proBDNF and BDNF ELISA experiment.

### BDNF and proBDNF ELISA

BDNF and proBDNF protein levels were determined by a two-side ELISA (Genstar, China, C643-02 and C546-02) as described by the manufacturer. Plasma were loaded directly into 96 well plates without dilution. Absorbance was recorded and analyzed using a Cytation BioTeK plate reader (Winooski, USA). BDNF and proBDNF concentration (pg/ml) was normalized to the volume of plasma in each sample.

## Statistics

Typically, statistical significance was determined using Student’s t-tests after confirming normal distribution of the data. In multi-bar figures, statistical significance was determined by analysis of variance followed by a post hoc test. Data are represented as the mean ± SEM.

## Acknowledgements

This work was supported by the Grant-in-Aid for Scientific Research on Priority Areas-Elucidation of neural network function in the brain-from the Ministry of Education, Culture, Sports, Science and Technology of Japan (40344171) (M.K.), by JST, CREST (T.M., H.K., T.H. and M.K.), and by the National Natural Science Foundation of China (31730034, 81501105), Beijing Municipal Science & Technology Commission (Z151100003915118), and Shenzhen Science, Technology and Innovation Commission (JCYJ20170411152419928 and (GJHZ20170314151528005) to B.L.. The funding organizations had no role in the study design, data collection and analysis, the decision to publish, or the preparation of the manuscript.

## Author Contributions

T.M., H.K., T.H., and M.K. (AIST) performed the biochemical assays, analyzed the data; T.M. and K.K.^a^ (AIST), performed the electrophysiological experiments and analyzed the data; K.T., T.M.^a,c^, K.K.^a,c^, M.O., and M.K. (AIST) engaged in the behavioral analysis of mice; T.M., K.T., M.O., M.K. performed the analysis of neuronal morphology and its quantitative analysis; T.B. and K.X. collected plasma samples of ASD patients; H.Y. conducted the ELISA analysis; H.Y., T.M., K.T., B.L., T.M., H.K., and M.K. wrote the paper; B.L., H.Y. and M.K. conceived and supervised the project.

---

**Figure 3-figure supplement 1.**
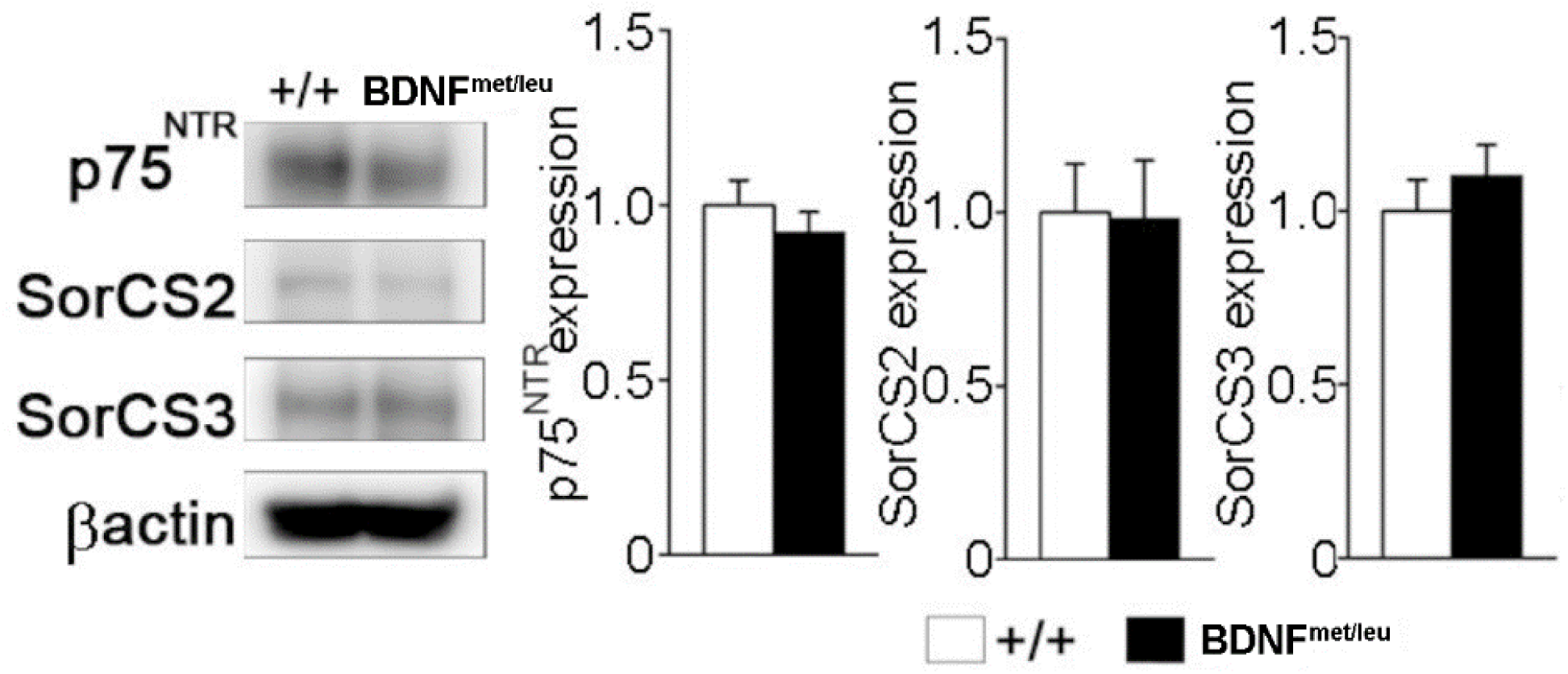
Immunoblot analyses of p75^NTR^, SorCS2, and SorCS3 expression in hippocampal lysates (25 μg of protein) from BDNF^+/+^ and BDNF^met/leu^ mice. The expression level of indicated proteins was normalized to that of beta-actin. The left panel shows a representative blot and the right panel shows quantification of the data (p75NTR, p = 0.18; SorCS2, p = 0.46; SorCS3, p = 0.77, by Student’s t-test.). N = 3 mice per group.

**Figure 5-figure supplement 1.**
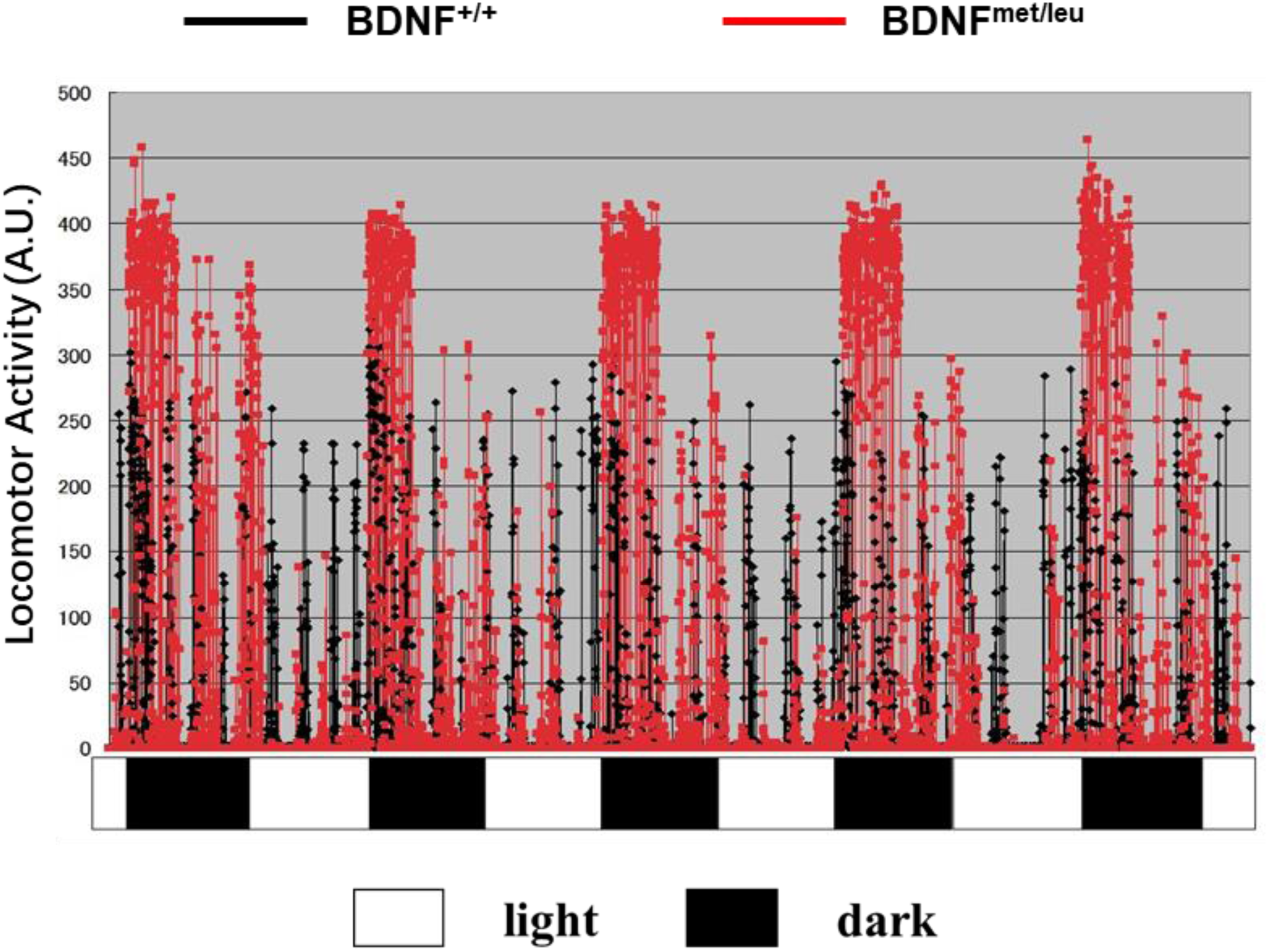
The recording of locomotor activity of BDNF^+/+^ (N=4) and BDNF^met/leu^ (N=4) mice during 4.5 days, with 12 h light and 12 h dark phases. Y axis indicates the horizontal activity for 5 minutes.

